# Leveraging ancestral recombination graphs for quantitative genetic analysis of rice yield in *indica* and *japonica* subspecies

**DOI:** 10.1101/2025.01.14.633033

**Authors:** Inés Rebollo, Daniel Tolhurst, Jana Obšteter, Juan Rosas, Gregor Gorjanc

**Affiliations:** Instituto Nacional de Investigación Agropecuaria (INIA), Estación Experimental Las Brujas, Ruta 48 km 10, Canelones, Uruguay; Department of Statistics, Universidad de la República, College of Agriculture, Garzón 780, Montevideo, Uruguay; The Roslin Institute and Royal (Dick) School of Veterinary Medicine, The University of Edinburgh, Edinburgh, UK; Agricultural Institute of Slovenia, Ljubljana, Slovenia

**Keywords:** Ancestral Recombination Graph, Genomic prediction, Rice

## Abstract

*Rice (Oryza sativa* L.) has two main subspecies, *indica* and *japonica*, which coexist in many regions but are often treated separately during breeding. Combining both subspecies in quantitative genetic analyses could enhance genetic improvement, however, this requires appropriately modelling their genetic history. The ancestral recombination graph (ARG) is an effective population genetics tool that comprehensively and succinctly represents a species’ genetic history. This study evaluated the use of an ARG, encoded as a tree sequence, to improve quantitative genetic analyses of *indica* and *japonica* rice. Using data from Uruguay’s National Rice Breeding Program, we inferred ancestral alleles, constructed and dated an ARG, and examined its application in genomic prediction and genome-wide association studies. We compared the predictive ability of a branch-based relationship matrix (BRM) built from an ARG against conventional relationship matrices from pedigree and single nucleotide polymorphism (SNP) site data. We then estimated BRM’s SNP site effects to identify potential sites of interest and better understand how these map onto the tree sequence branches. The results showed that the ARG captured key biological signals, encoded genomic data more efficiently than conventional formats, and resulted in the highest predictive ability when combining both subspecies. Although the ARG-based approach did not substantially outperform conventional approaches for between-species prediction, this approach holds promise for plant breeding with larger datasets and could enhance genome-wide association studies by elucidating haplotype ancestry and the evolution of their value. Overall, our results demonstrated the potential of ARGs for the quantitative genetic analysis of diverse populations.

## Introduction

Rice (*Oryza sativa* L.) is a major food staple, with two main subspecies, *indica* and *japonica*. Despite their co-existence in many regions, *indica* and *japonica* are generally treated as separate populations in breeding programs. Combining these populations could improve the efficacy of current genomic prediction and genome-wide association studies; however, this requires appropriate modelling of the populations’ genetic history. Ancestral recombination graph (ARG) is the ultimate way to represent the complete genetic history of individuals and their genomes. ARGs have been shown to be effective in many population genetic studies, and hold great potential to improve quantitative genetic modelling. In this study, we evaluate the use of an ARG, encoded as tree sequences, to improve quantitative genetic modelling in *indica* and *japonica* rice subspecies.

Genetic analysis of cultivated rice varieties revealed two main subspecies, *indica* and *japonica* (Glaszmann 1987; Garris *et al*. 2005), which share a common ancestor with the wild *O. rufipogon* species (Huang *et al*. 2012). They were estimated to have diverged ∼0.4 million years ago (Zhu and Ge 2005), by a single domestication event with subsequent divergence and migration between subspecies, or by multiple domestication events (Caicedo *et al*. 2007; Liu *et al*. 2024). Due to this deep divergence, the cultivation, breeding, and quantitative genetic analyses are generally conducted separately for *indica* and *japonica*. However, in some countries in Asia and elsewhere, such as Uruguay in South America, *indica* and *japonica* coexist (Martínez *et al*. 2014). In these places, small- and medium-scale rice breeding programs of *indica* and *japonica* occur, and individual populations of each subspecies have limited size. This raises the possibility of considering joint quantitative genetic analyses and breeding operations for both subspecies, but the genetic history of rice subspecies requires appropriate modelling (Caicedo *et al*. 2007; Liu *et al*. 2024).

Rice breeders generally select and improve their populations based on estimated genetic values, which are obtained from a linear mixed model with a relationship matrix (RM) describing the genetic similarity between individuals (Henderson 1984). The RM can be built from different sources, such as pedigree or genomic data. A pedigree-based relationship matrix (PRM) expresses the expected identity by descent between pairs of individuals relative to the base population of a known pedigree (Henderson 1984; Mrode 2014). In contrast, the genomic RMs proposed by VanRaden (2008) express the realized identity by state across single nucleotide polymorphism (SNP) sites. This notion captures the relationships between pairs of individuals due to shared mutations that segregate between and within families. We refer to this type of RM as a site-based relationship matrix (SRM), because it is based on current variation at polymorphic sites typed by SNP markers. The SRM enhances quantitative genetic modelling in terms of genomic prediction relative to the PRM and enables genome-wide association studies (de los Campos *et al*. 2013; Yu *et al*. 2006). However, the commonly used SRM does not fully reflect the genetic history of sampled individuals because it does not capture all SNP sites, and origin of mutations (Young 2022; Fan *et al*. 2022). This can reduce the efficacy of these applications for multiple (divergent) populations, like *indica* and *japonica* rice, so there remains the need to consider appropriate RMs for joint quantitative genetic analyses of these subspecies.

Joint quantitative genetic analysis of multiple populations is an active area of research (Martin *et al*. 2017; Ding *et al*. 2023; Mester *et al*. 2023; Warburton *et al*. 2023; Zhang *et al*. 2023b; Sun *et al*. 2024). One of the key considerations for joint analyses of small populations is that the construction of a sufficiently large training dataset is likely to include diverged populations with specific recombination and mutation events, which can counteract the advantages of constructing a larger training dataset (Lorenz and Smith 2015). Therefore, the joint quantitative genetic analysis of *indica* and *japonica* could be beneficial, but requires a better representation and modelling of the quantitative genetic diversity between and within the subspecies.

An ARG describes the genetic history of sampled genomes by encoding their shared ancestry, mutation, and recombination events (Griffiths and Marjoram 1996; Brandt *et al*. 2024; Lewanski *et al*. 2024; Nielsen *et al*. 2024; Wong *et al*. 2024). ARGs have been used to recover relationships between individuals and populations (e.g., Kelleher *et al*. 2019; Wohns *et al*. 2022; Fan *et al*. 2022), estimate coalescence times (e.g., Brandt *et al*. 2022), map quantitative trait loci (QTL) (e.g., Link *et al*. 2023), and estimate heritability and genome-wide association (e.g., Zhang *et al*. 2023a). Within an ARG, shared ancestry at a locus is encoded by a local tree, where the nodes represent the sampled haploid genomes and their ancestors (Brandt *et al*. 2024; Lewanski *et al*. 2024; Nielsen *et al*. 2024; Wong *et al*. 2024). The nodes are connected with edges, represented as branches in a local tree, which indicate the lineages of descent between haploid genomes. Mutations on the branches generate variation between genomes at a given locus. Recombinations between loci change the lines of descent between haploid genomes and hence change the topology of local trees along the genome (Brandt *et al*. 2024; Lewanski *et al*. 2024; Nielsen *et al*. 2024; Wong *et al*. 2024). Recent advances in methodology and software implementations now enable the inference of ARGs for many genomes and populations (Speidel *et al*. 2019; Kelleher *et al*. 2019; Zhang *et al*. 2023a; Harris 2023; Gunnarsson *et al*. 2024; Nielsen *et al*. 2024)

In this study, we use the tree sequence representation of an ARG to improve quantitative genetic modelling of *indica* and *japonica* rice subspecies. A tree sequence succinctly encodes a sequence of local trees along the genome (Kelleher *et al*. 2016, 2018). Tree sequences enable efficient genome storage in simulations (Kelleher *et al*. 2016, 2018; Haller *et al*. 2019) and population genetic studies (Kelleher *et al*. 2019), as well as fast calculations of population genetic statistics (Ralph *et al*. 2020). There is now a collection of software to work with tree sequences, such as inferring tree sequence from phased genomes (Kelleher *et al*. 2019), dating nodes of a tree sequence (Wohns *et al*. 2022), and calculating population genetic statistics from a tree sequence (Ralph *et al*. 2020). Importantly, the work of Ralph (2019) and Ralph *et al*. (2020) provide a framework for calculating population genetic statistics based on site and branch information, including a site-based and branch-based RM from an ARG (B. Lehmann et al., personal communication, November 27, 2020). We used tskit implementation of branch-based RM (Kelleher *et al*. 2021a), which we call BRM. The BRM captures the rich genetic history of sampled individuals, including untyped sites, linkage between sites, and enables estimation of haplotype effects. Recently, Fan *et al*. (2022), Zhang *et al*. (2023a), Link *et al*. (2023), Gunnarsson *et al*. (2024), and Zhu *et al*. (2024) showed that the BRM captures the genetic diversity better than the SRM, opening opportunities to improve quantitative genetic modelling across multiple populations, such as rice subspecies. This work aimed to evaluate the use of tree sequences as a tool to build a RM for a joint quantitative genetic analysis of *indica* and *japonica* rice genotypes from a breeding program in Uruguay. To this end, we inferred and analyzed tree sequences for a sample of individuals from the Uruguayan National Rice Breeding Program, built the BRM, and used it for genomic prediction and genome-wide association study of grain yield. The results of this study can guide strategies for joint selective breeding and management of genetic diversity in *indica* and *japonica* rice and other species with a similar population structure.

## Materials and methods

We used genomic, pedigree, and phenotypic data from the Uruguayan National Rice Breeding Program of *indica* and *japonica* rice between 1997 and 2020 (Martínez *et al*. 2014; Rebollo *et al*. 2023a). We first inferred the rice ancestral alleles and the ARG for all individuals using a tree sequence with dated nodes. We then analyzed the properties of inferred and dated tree sequences, including genealogical nearest neighbor analysis, and inspected the local trees at two selected regions. Using different cross-validation (CV) scenarios, we evaluated the ability of linear mixed models using either the PRM, SRM, or BRM to predict grain yield between and within the subspecies. Finally, we inspected estimated SNP site effects to understand how the estimates map onto branches of the ARG.

### Data

Genomic, pedigree, and phenotypic data were retrieved from the Uruguayan National Rice Breeding Program. A detailed description of the data and their availability is given in Rebollo *et al*. (2023a), and summarised in the following.

Genomic data were available for 965 late-stage inbred lines of *indica* (395 individuals) and *japonica* (570 individuals), generated with Genotyping-by-Sequencing (GBS) of ApeKI enzyme-cut DNA fragments (Elshire *et al*. 2011; Rebollo *et al*. 2023a). We aligned the sequences to the Nipponbare reference genome MSU version 7.0 (Kawahara *et al*. 2013) using bwa (Li and Durbin 2009). Next, we used the TASSEL 5.0 pipeline (Bradbury *et al*. 2007) to call SNP sites and individual genotypes. We performed quality control and retained SNP sites with minor allele frequency ≥0.03, missing data ≤ 50%, and observed heterozygosity ≤15%. The resulting genotypes were phased and imputed with BEAGLE version 5.1 (Browning *et al*. 2018). We performed all steps above for both subspecies together (ALL), and separately for *indica* (IND) and *japonica* (JAP). A total of 61,260 SNP sites were retained for ALL, 50,854 for IND, and 23,614 for JAP. Some SNP sites were shared between genomic datasets, while some were exclusive to each dataset (Supplementary Figure 1). In line with the aims of the study, the majority of the analyses and subsequent results focus on the ALL genomic dataset.

Pedigree data were available for 915 of the 965 individuals with genomic data (95%), producing a total of 1,207 individuals in the pedigree including 292 ancestors. There were up to six generations of ancestors with 204 founder individuals. Individuals without pedigree data were included as nominally unrelated to the remaining individuals, i.e., treated as founders. The number of self-pollination generations were stored with the pedigree for later calculation of the PRM.

Grain yield data were available for 936 of the 965 individuals with genomic data (97%), which are a subset of the 19,447 individuals tested across 23 years of field evaluation (Rebollo *et al*. 2023a). The selected grain yield data came from 846 field trials conducted between 1997 and 2020 in three locations: Paso de la Laguna (33.27 S, 54.17 W) with 743 trials; Paso Farías (30.54 S, 57.26 W) with 92 trials; and Pueblo del Barro (31.93 S, 55.38 W) with 11 trials. Each trial had a randomized complete block design with two to four blocks. All trials were conducted under current production management standards for irrigated conditions (Batello *et al*. 2013). Grain yield was recorded from 1.20 m x 2.0 m plots (2.4 m^2^) and converted to kg/ha. The total number of recorded plots was 23,311. The number of individuals present each year ranged from 12 to 653 (mean of 173), while the number of individuals shared between pairs of years ranged from 3 to 640. Field trials were grouped into 49 environments, defined by their year location combination. Each environment had 1 to 48 field trials (mean of 17), and 10 to 653 individuals with genomic data (mean of 95). Environments were excluded if they contained less than 20 individuals. The final genomic, pedigree, and phenotypic datasets included 22,741 records across 40 environments for 936 individuals, with 381 *indica* and 555 *japonica* individuals.

### Ancestral recombination graph

We combined our genomic dataset and Ensembl’s alignment of *Oryza sativa* with other *Oryza spp*. (Yates *et al*. 2021) to infer the rice ancestral alleles using *O. rufipogon, O. glaberrima*, and *O. meridionalis* as external species. We converted Ensembl Multiple Format (EMF) to Multiple Alignment Format (MAF) and accessed specific regions in the external species corresponding to SNP sites in our genomic dataset using WGAbed (Corcoran and Barton 2021). We inferred the rice ancestral alleles using est-sfs (Keightley and Jackson 2018) with Rate-6 model of nucleotide substitution and 10 random starting values. We declared the most probable allele at the common ancestor with the first out-group (*O. rufipogon*) as the ancestral allele. Where we could not infer the ancestral allele, we declared the major allele as ancestral. We summarised this information by computing the total number of SNP sites inferred, and the number and percentage of times that the ancestral allele was the major, minor, or another allele. Finally, we computed and compared the alternative (i.e., non-ancestral) and minor allele frequency spectrum of the ALL, IND, and JAP genomic datasets.

We inferred an ARG encoded as a tree sequence for each chromosome with tsinfer 0.2.1 (Kelleher *et al*. 2019, 2021b). The succinct encoding of tree sequences is achieved by a collection of core tables that describe: (i) haploid genomes (nodes) and their date of existence, (ii) shared DNA between nodes (edges/branches), (iii) SNP sites with their ancestral state, and (iv) mutation events on the edges giving rise to polymorphisms at the sites, alleles of the nodes, and genotypes of individuals. For inferring tree sequences, we used a recombination rate of 4.53 *×* 10^−8^ per base pair per generation (Si *et al*. 2015), the default mismatch ratio parameter of 1, and physical SNP positions from the Variant Call Format (VCF) file. Finally, we summarised information on the number of nodes, edges, trees, SNP sites, and mutations for the tree sequence of each chromosome.

We dated the ARG by inferring the age of the nodes with tsdate 0.1.4 (Wohns *et al*. 2022). For this inference, we used a mutation rate of 2.2 *×* 10^−9^ per base pair per generation (Yang *et al*. 2015). This dating required knowledge of the effective population size (*Ne*) and as there are contrasting estimates of *Ne* depending on the type of data and time, we used three different values (23, 1,500, and 150,000) to evaluate its impact on dating and downstream analyses. We estimated the current *Ne* of 23 from the pedigree-based rate of coancestry (Pérez-Enciso 1995), which is concordant with Rutkoski (2019). We estimated the recent *Ne* of 1500 from the linkage disequilibrium of the ALL genomic dataset at 50 generations ago with GONE 29.08.2021 (Santiago *et al*. 2020). We ran GONE with default parameters, except for the recombination rate mentioned above and the maximum number of recombination bins set to 0.005. We used the ancient *Ne* of 150,000 from the work of Caicedo *et al*. (2007). Results presented in the main text correspond to *Ne* = 1500 and we show the sensitivity of the results to the other two *Ne* values in Supplementary Material. From the dated ARG, we estimated the age of SNP sites as the age of the first ancestor above which we inferred a mutation. Finally, we summarized the information on the age of ancestors, mutations, and SNP sites.

To extract biological signals from the ARG, we calculated the genealogical nearest neighbors (GNN) statistic (Kelleher *et al*. 2019) for all individuals and inspected two local trees of interest. The GNN describes the topology of an ARG with respect to a reference set (subspecies), to describe the identity of nearest neighbors within a local tree and summarized across the local trees (Kelleher *et al*. 2019). We calculated a GNN matrix for each chromosome between all pairs of individuals with tskit 0.3.7. We then combined these matrices into a single GNN matrix by summing across chromosomes and weighting by their relative proportions of the genome. We hierarchically clustered the individuals with the average method using SciPy (Virtanen *et al*. 2020). The first local tree of interest was between positions 32,645,584 and 32,648,666 bp of chromosome three, covering the drought and salt tolerance (*DST*) gene, which is associated with panicle length in *japonica* only (Bai *et al*. 2016). The second local tree of interest was between positions 31,583,546 and 31,596,967 bp of chromosome four, covering position 31,590,530 bp of chromosome four which is associated with the number of panicle secondary branches (NSB) and number of secondary spikelets per secondary branch (NSSB) in both *indica* and *japonica* (Bai *et al*. 2016).

### Relationship matrices

We calculated the PRM considering the number of generations of self-pollination with preGSf90 (Aguilar *et al*. 2014; Rebollo *et al*. 2020). The SRM was calculated with the rrBLUP R package (Endelman 2011) as **MM**^⊤^/*c*, where **M** is the SNP genotype matrix centered by the mean allele dosage at each site and *c* is the sum of the expected heterozygosities across SNP sites (VanRaden 2008). A BRM was calculated for each chromosome from the inferred tree sequence by computing the total area of shared branches between and within each pair of homologous chromosomes of all individuals, relative to the remaining homologous chromosome pairs using tskit 0.3.7 (Kelleher *et al*. 2021a) (B. Lehmann et al., personal communication, November 27, 2020). These matrices were then combined into a single BRM by summing across chromosomes and weighting by their relative proportions of the genome. Branch-based statistics, such as the BRM, are calculated from the length and span of branches, giving the expected value of its dual site-based statistic under the infinite-sites mutation model (Ralph 2019; Ralph *et al*. 2020). For the purpose of subsequent quantitative genetic analyses, we write the BRM as **MDM**^⊤^, where **M** is the SNP genotype matrix and **D** is a relationship matrix between SNP site effects, induced through the tree sequence structure (see the Appendix).

### Quantitative genetic analysis

We fitted various linear mixed models, with a particular focus on modelling different structures for genotype-by-environment (GxE) interaction and different genotype relationship matrices. The phenotypic dataset comprised *v* = 936 individuals evaluated in *t* = 828 trials across *p* = 40 environments (year locations) with *n* = 22, 741 records in total. Let the *n*-vector of phenotypic records be given by 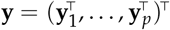, where **y**_*j*_ is the *n*_*j*_-vector for the *j*^*th*^ environment. The linear mixed model for **y** is given by:

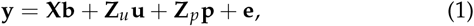

where **b** is a vector of fixed effects with design matrix **X, u** is the *vp*-vector of random genetic values, ordered as individuals in environments, with design matrix **Z**_*u*_, **p** is a vector of random non-genetic effects with design matrix **Z**_*p*_, and **e** is the *n*-vector of residuals. The fixed effects included the overall and environment means, while the random non-genetic effects captured the blocking structure of trials within environments. The random effects and residuals were assumed to be mutually independent following a multivariate normal distribution.

We fitted seven linear mixed models, as summarised in Supplementary Table 1. All models included diagonal covariance structures with separate block and residual variances for each environment. Note that the absence of field layouts precluded fitting residual spatial models. All models also included a separable covariance structure for the genetic values given by var(**u**) = **G**_*e*_ ⊗ **G**, where **G**_*e*_ is an unknown covariance structure between environments and **G** is a known RM between individuals. We considered seven forms of **G**_*e*_ to model different GxE interaction patterns and three forms of **G** to model the different sources of genetic data (i.e., the PRM, SRM, and BRM). Details on the different forms of **G**_*e*_ can be found in Tolhurst *et al*. (2022), but briefly:

- Model 1 assumes independent genetic values between environments, modelled by a diagonal covariance structure with a separate genetic variance for each environment.
- Model 2 assumes correlated genetic values between environments, modelled by a uniform covariance structure with a single genetic variance and covariance.
- Model 3 is an extension of model 2 that includes an additional interaction variance across environments, producing a compound symmetric covariance structure.
- Model 4 is an extension of model 3 that includes a separate interaction variance for each environment, rather than a single variance across environments.
- Models 5, 6, and 7 include factor analytic covariance structures of order 1, 2, and 3, respectively. These models fit a different genetic variance for each environment and a different genetic covariance for each pair of environments.

Model 2 comprises a single mean genetic value across environments for each genotype while the remaining models comprise specific genetic values for each genotype by environment combination. All models were fitted using ASReml-R (Butler *et al*. 2017), which obtains Residual Maximum Likelihood (REML) estimates of the variance parameters and empirical Best Linear Unbiased Predictions (BLUPs) of the random effects. Model fit was assessed using the residual log-likelihood, AIC, and percentage of variance explained (*v*_*e*_). The *v*_*e*_ was calculated as the variance explained by the correlated genetic values as a percentage of the total genetic variance following Tolhurst *et al*. (2019). Models 2 and 4 were then used for CV and genome-wide association, as described in the following.

We investigated how the different RMs impact the predictive ability of the different linear mixed models. Due to generational overlap in our data and a low number of individuals in the last generations, we could not perform forward validation of the genomic prediction of grain yield and have performed a five-fold CV instead. To ensure a sufficiently structured and powered CV, we used a subset of the phenotypic records corresponding to the three years with the highest number of shared individuals (2011-2013). The subset comprised 93 trials at Paso de la Laguna with 327 *indica* and 330 *japonica* individuals in total. We used five-fold CV, with the dataset partitioned into five sets containing about 65 *indica* and 66 *japonica* individuals. Each set was used as a validation set while the remaining four sets were used to train the model for prediction. We tested five CV scenarios: (i)*CV*_*IJ* →*I J*_ trained on and predicted both *indica* and *japonica*, (ii)*CV*_*I* →*J*_ trained on *indica* and predicted *japonica*, (iii) *CV*_*J* →*I*_ trained on *japonica* and predicted *indica*, (iv) *CV*_*I* →*I*_ trained on and predicted *indica*, and (v) *CV*_*J* →*J*_ trained on and predicted *japonica*. Each scenario was replicated 1,000 times. We computed the predictive ability for each scenario as Pearson’s correlation coefficient (*r*) between the predicted mean genetic values and mean phenotypic values across environments. For model 4, *r* was also computed between the predicted specific genetic values and mean phenotypic values within environments.

We performed a genome-wide association study (GWAS) on the complete data set, with standardised effects of the SNP sites obtained following Gualdrón Duarte *et al*. (2014):

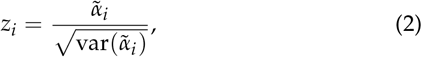

where 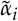 is the the BLUP of the *i*^*th*^ SNP site effect and 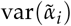 is its variance. These components were obtained from the genetic values parameterised by the SRM and BRM following Tolhurst *et al*. (2019), as described in the Appendix. Note, however, since the matrix **D** was not readily available for this study we were limited to obtaining near-unique BLUPs and variances for the BRM. The associated *p*-values from both RMs were displayed using Manhattan plots, with significance assessed at *p* = 0.05 using Li and Ji (2005) correction that accounts for the effective number of independent tests. For each model, we also calculated Pearson’s correlation coefficient between the predicted SNP site effects obtained with each RM.

Finally, we selected a region spanning two GWAS hits and examined how the effects of the SNP sites from model 2 with the BRM mapped to the branches of the tree sequence and corresponding haplotype effects. A haplotype was defined as the combination of the allelic states of contiguous SNP sites. Haplotype effects were estimated as a linear combination of their SNP site effects 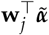 and standardised using:

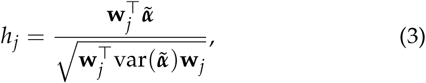

where **w**_*j*_ denotes a vector of 0s and 1s for allelic states for the *j*^*th*^ haplotype and 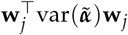 the variance of the haplotype effect, with 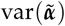 being the variance-covariance matrix of 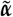. Note that our interpretation of the *h*_*j*_ is limited due to the near-unique solutions of the 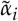 mentioned above.

## Results

Overall, our results show that: (i) tree sequences captured the underlying biological signal from the genomic data; (ii) tree sequences encoded the genomic data more succinctly than the standard format; (iii) the site-based (SRM) and branch-based (BRM) relationship matrices were highly correlated, as were subsequent grain yield predictions; (iv) the BRM achieved the highest predictive ability when information from both *indica* and *japonica* was analyzed; and (iv) SNP site effects mapped onto branches of an ARG provide a way to estimate haplotype effects and give insight into the evolution of their value.

The results are structured as follows. We initially demonstrate the usefulness of tree sequences to capture biological signal summarised with genealogical nearest neighbors (GNN) and local trees for regions of interest. We then summarise the ancestral alleles and tree sequences inference. The latter includes the summary of memory usage and compression, as well as the estimated age of ancestors, mutations, and SNP sites. Finally, we present the results of quantitative genetic analyses based on tree sequences, including genomic prediction and GWAS. Here, we present the results for *Ne* = 1,500 when dating the tree sequences. The results for *Ne* = 23 and 150,000 are presented in the Supplementary Material.

Our results show that tree sequences captured the underlying biological signal in the genomic data. The GNN captured the inherent population structure and clearly separated the *indica* and *japonica* rice subspecies (Figure 1). Within subspecies, the GNN revealed a more pronounced structure for *indica* compared to *japonica*, possibly indicating a stronger family structure. Figure 1 also demonstrated admixed individuals between subspecies or potentially mislabeled individuals. We found a differential local structure by inspecting two interesting regions of the genome (Figure 2). Both regions had two SNP sites and two local trees (one SNP per tree), but the adjacent local trees were practically indistinguishable. The trees corresponding to the first SNP for both regions are shown in Figure 2 while the trees of the second SNP are shown in Supplementary Figure 2. The tree spanning the *DST* gene showed a clear and deep separation between *indica* and *japonica* (Figure 2A). Here, 98% of *japonica* individuals had the ancestral haplotype at both SNP sites, while 77% of *indica* individuals had a mutation at the first SNP and 19% at the second SNP (Supplementary Table 3). The tree covering the locus associated with NSB and NSSB showed that this region was segregating in both *indica* and *japonica* (Figure 2B). Here, 27% of *indica* individuals and 80% of *japonica* individuals had the ancestral haplotype at both SNP sites while 71% of *indica* individuals had a mutation at the second SNP and 19% of *japonica* individuals had a mutation at the first SNP (Supplementary Table 3).

**Figure 1.**
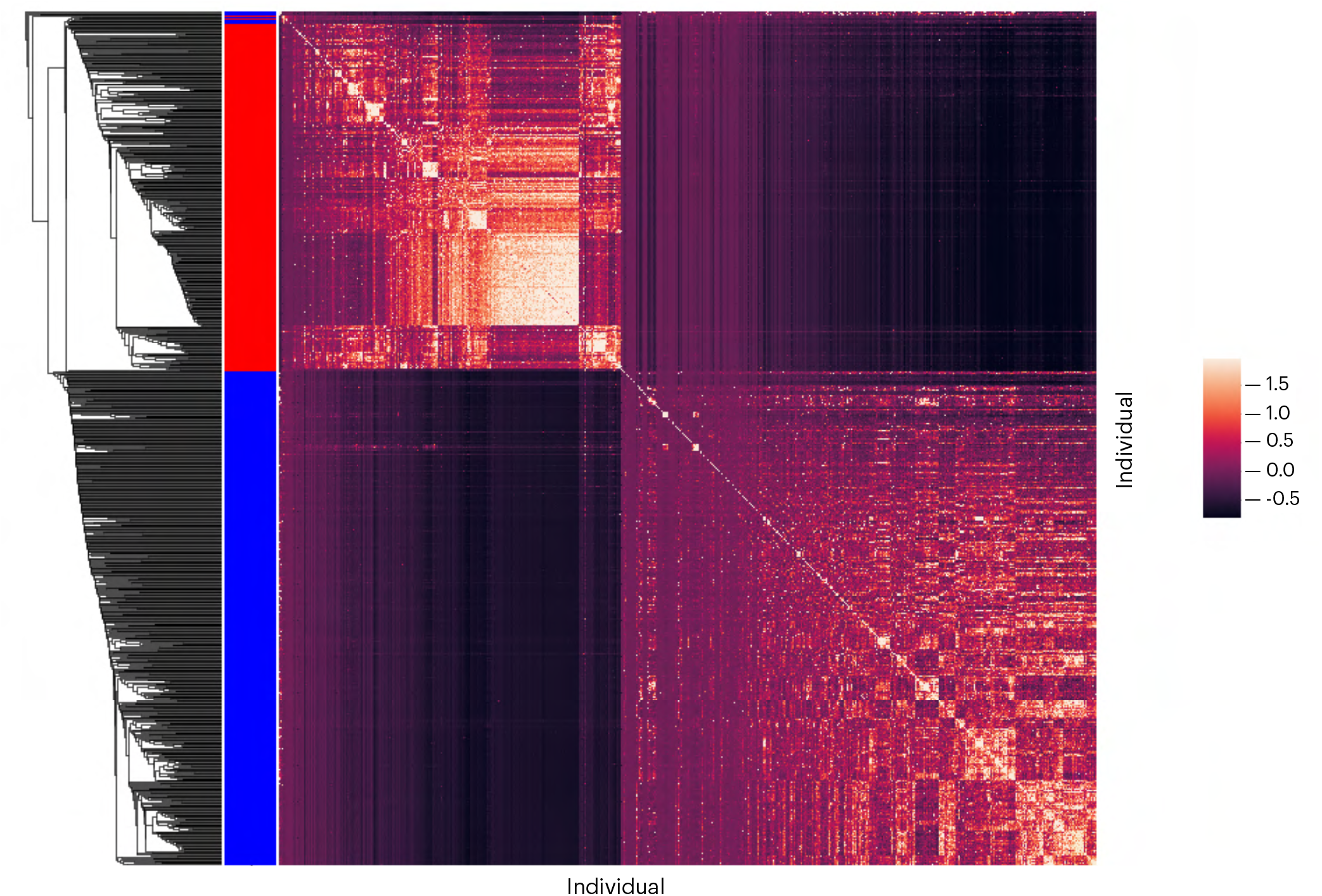
Genealogical nearest neighbors (GNN) of *indica* individuals in red and *japonica* individuals in blue. The GNN matrix was hierarchically clustered using the average method, shows the inherent population structure, and clearly separated the *indica* and *japonica* rice subspecies.

**Figure 2.**
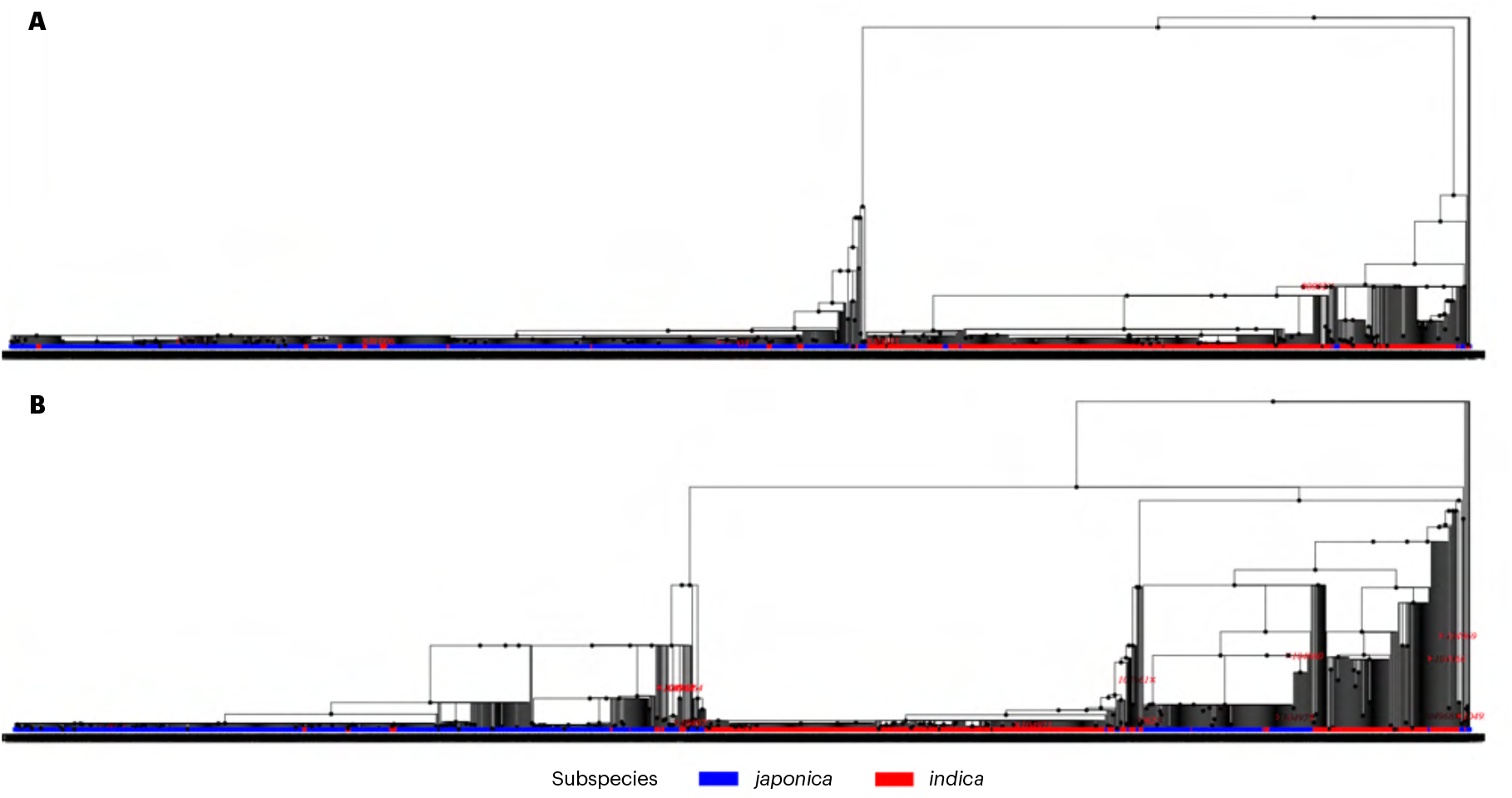
Differential local tree structure at genome positions: (A) the Drought and Salt Tolerance (*DST*) locus associated with panicle length only in *japonica*; and (B) a locus associated with number of panicle secondary branches and number of spikelets per panicle secondary branch in both *indica* and *japonica*. The tree in (A) shows a very clear and deep separation between *indica* and *japonica* while the tree in (B) shows segregation in both *indica* and *japonica*.

We inferred ancestral alleles for about 80% of the loci in the three genomic datasets (ALL, IND, and JAP; Table 1). Ancestral alleles matched the major allele at 68% loci in the ALL dataset, 59% loci in the JAP dataset, and 52% loci in the IND dataset. In less than 1% of all cases, the ancestral allele was neither the major nor the minor allele. Some alleles were more common in *indica* while others were more common in *japonica*. The distribution of the alternative (non-ancestral) allele frequency within each subspecies was U-shaped, with an excess of the lower alternative allele frequency for *japonica* (Figure 3A). However, when combined, the distribution indicated differences in alternative allele frequencies between the two subspecies (Figure 3B). Some SNP sites had a low frequency in one subspecies and a high frequency in the other, and vice-versa (Figure 3C). Similar results were demonstrated by the distribution of minor allele frequencies (Supplementary Figure 3).

**Table 1.** Characterization of the ancestral allele inference for the genomic datasets with all individuals from both subspecies (ALL) and from each subspecies separately (IND for *indica* and JAP for *japonica*). Percentages of the total number of ancestral alleles inferred are presented in parentheses.

| Genomic dataset | Total sites | Alleles inferred | Ancestral = Major | Ancestral = Minor | Ancestral ≠ Minor ≠ Major |
| --- | --- | --- | --- | --- | --- |
| ALL | 61,260 | 49,518 (80.8%) | 33,485 (67.6%) | 15,802 (31.9%) | 231 (0.5%) |
| IND | 50,854 | 40,891 (80.4%) | 21,253 (52.0%) | 19,363 (47.3%) | 275 (0.7%) |
| JAP | 23,614 | 18,519 (78.4%) | 10,905 (58.9%) | 7,511 (40.6%) | 94 (0.5%) |

**Figure 3.**
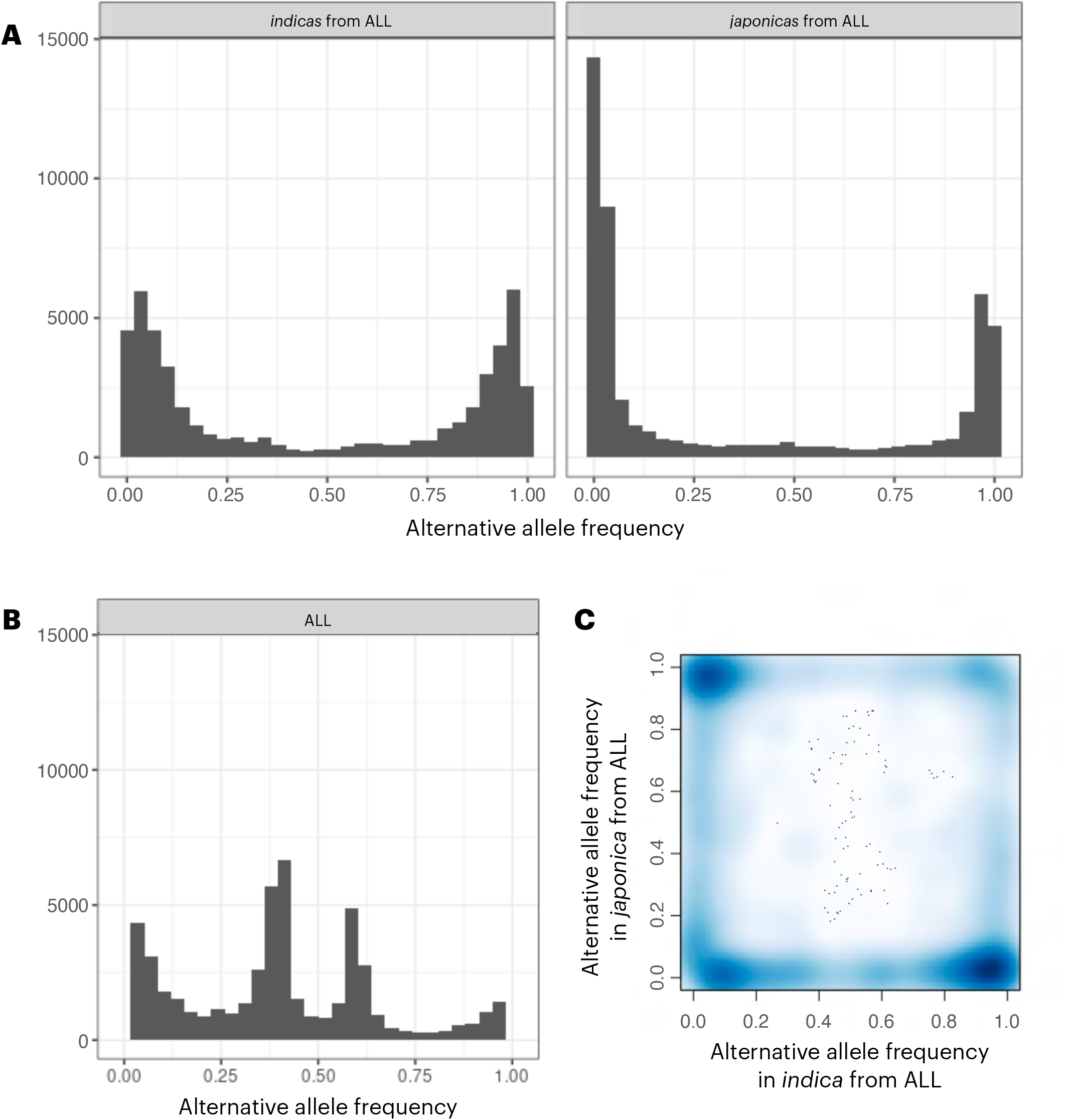
Alternative (non-ancestral) allele frequency spectrum: (A) alternative allele frequency of the subset of *indica* and *japonica* from the genomic dataset with all individuals (ALL); (B) alternative allele frequency of the ALL genomic dataset; and (C) scatter plot between the alternative allele frequency of *indica* and *japonica* from the ALL genomic dataset. The histogram in (a) shows a U-shape distribution for each subspecies, as expected for neutral diversity, with an excess of the lower alternative allele frequency for *japonica*.

The inferred tree sequences encoded the genomic data more succinctly than the standard format. The combined tree sequence across chromosomes included 95,612 nodes, 693,524 edges, 31,925 local trees, 61,260 SNP sites, and 1,031,672 mutations in total (Table 2). These statistics varied for individual chromosomes, and were generally well-correlated with the chromosome length. The combined file size of the dated trees across chromosomes was 62 Mb while the standard VCF file was 228 Mb, i.e., almost four times larger.

**Table 2.** Number of nodes, edges, trees, sites, and mutations in total for the combined tree sequence and separately for each chromo-some.

| Chromosome | Nodes | Edges | Trees | Sites | Mutations |
| --- | --- | --- | --- | --- | --- |
| TOTAL | 95,612 | 693,524 | 31,925 | 61,260 | 1,031,672 |
| 1 | 11,195 | 73,825 | 4,429 | 8,245 | 119,709 |
| 2 | 9,584 | 67,879 | 3,543 | 6,549 | 106,885 |
| 3 | 8,568 | 55,794 | 3,181 | 6,126 | 91,245 |
| 4 | 9,647 | 81,314 | 3,195 | 6,198 | 116,737 |
| 5 | 6,159 | 45,476 | 1,952 | 3,815 | 69,731 |
| 6 | 8,228 | 58,815 | 2,719 | 5,442 | 82,933 |
| 7 | 6,776 | 49,100 | 2,204 | 4,239 | 63,656 |
| 8 | 7,572 | 58,206 | 2,428 | 4,577 | 79,862 |
| 9 | 6,620 | 46,421 | 1,971 | 3,977 | 80,205 |
| 10 | 6,513 | 45,489 | 1,930 | 3,718 | 77,165 |
| 11 | 8,671 | 70,554 | 2,631 | 4,988 | 86,870 |
| 12 | 6,079 | 40,651 | 1,742 | 3,386 | 56,674 |

The age distributions for nodes (ancestors), mutations, and SNP sites (i.e., first mutation at each site) were heavily right-skewed toward the present (Figure 4). For the nodes, this is expected due to the coalescence of nodes backward in time. Furthermore, since the dataset involved a breeding population, we expect to observe rapid coalescence due to selection and a small number of parents. Assuming *Ne* = 1,500, 70.4% of nodes were estimated to be younger than 1,000 generations, 61.5% younger than 500 generations, and 41.2% younger than 100 generations. Conversely, 54.8% of mutations were estimated to be younger than 2,500 generations, 30.4% were estimated between 2,500 and 5,000 generations, and 14.8% were estimated above 5,000 generations. All chromosomes showed a similar age pattern for nodes, SNP sites, and mutations. The *Ne* used when dating the tree sequences strongly affected the estimated age of nodes, SNP sites, and mutations (Supplementary Figure 4).

**Figure 4.**
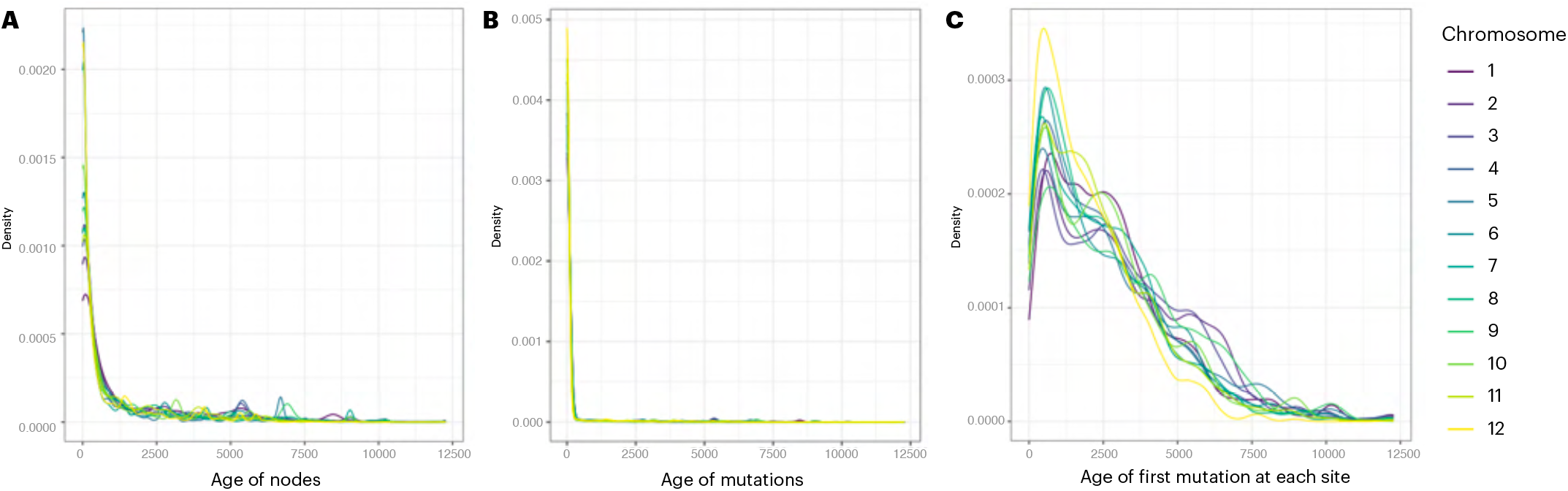
Tree sequence inference for each chromosome: (A) estimated age of nodes; (B) estimated age of mutations; and (C) estimated age of first mutation at each site. The ages in (A) represent the age of ancestors in the trees while the ages in (C) represent the age at which sites became polymorphic, i.e., the age of the oldest ancestor above which we inferred a mutation. All ages were heavily right-skewed toward the present, with all chromosomes presenting a similar pattern.

Heatmaps of the SRM and BRM revealed a very similar population structure with elements of the matrices being highly correlated, although on a different scale (Figure 5). The correlation between the diagonal elements of the SRM and BRM was 0.98 across both subspecies (0.75 for *indica* and 0.89 for *japonica*) while the correlation between the off-diagonal elements was 0.99 (1.00 for *indica* and 0.98 for *japonica*. The *Ne* used when dating the tree sequences impacted the resulting BRM; with *Ne* = 23 producing significantly lower values (Supplementary Figure 5).

**Figure 5.**
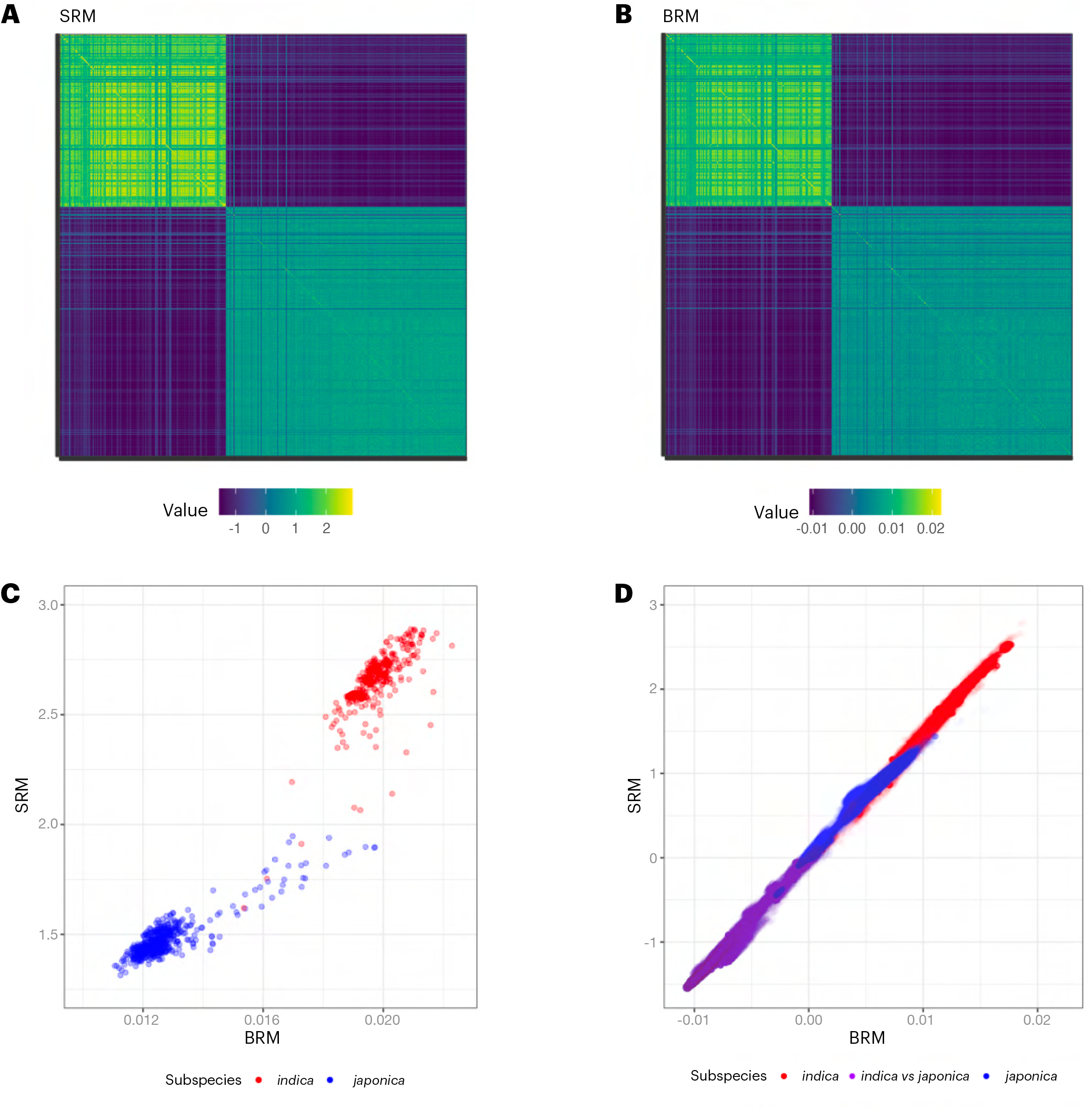
Comparison of the site-based (SRM) and branch-based (BRM) relationship matrices: (A) heatmap of the SRM; (B) heatmap of the BRM; (C) comparison of the diagonal elements of each matrix; and (D) comparison of the off-diagonal elements. The figure shows that the SRM and BRM revealed a very similar population structure with elements of the matrices being highly correlated, although on a different scale.

The predicted mean genetic values were highly correlated between the SRM and BRM when the subspecies in the training and prediction sets were the same (*r* = 0.96 − 0.99; Table 3). However, the correlations were lower when the subspecies in the training and prediction sets were different (*r* = 0.74 − 0.88). The equivalent correlations involving the PRM were much more varied, e.g., the correlation was low (*CV*_*I*→*J*_) and negative (*CV*_*J*→*I*_) when the training and prediction sets were different. Finally, note that the correlations involving the PRM were higher with the BRM than with the SRM.

**Table 3.** Correlation between predicted mean genetic values from model 2 and between predicted specific genetic values from model 4 for different relationship matrices and cross-validation scenarios. Presented is the mean correlation across 1000 replicates of each scenario, with standard deviations in parentheses.

| Scenario | Relationship matrix | Model 2 |  | Model 4 |  |
| --- | --- | --- | --- | --- | --- |
|  |  | SRM | BRM | SRM | BRM |
| $CV_{IJ \rightarrow IJ}$ | PRM | 0.78 (0.05) | 0.82 (0.05) | 0.74 (0.05) | 0.76 (0.05) |
|  | SRM |  | 0.98 (0.01) |  | 0.98 (0.01) |
|  | PRM | 0.70 (0.08) | 0.72 (0.07) | 0.70 (0.06) | 0.73 (0.05) |
|  | SRM |  | 0.97 (0.01) |  | 0.97 (0.01) |
| $CV_{I \rightarrow J}$ | PRM | 0.10 (0.15) | 0.13 (0.15) | 0.16 (0.12) | 0.20 (0.12) |
|  | SRM |  | 0.88 (0.05) |  | 0.84 (0.07) |
| $CV_{J \rightarrow I}$ | PRM | -0.16 (0.19) | -0.04 (0.19) | -0.08 (0.15) | 0.04 (0.13) |
|  | SRM |  | 0.74 (0.14) |  | 0.80 (0.15) |
| $CV_{I \rightarrow I}$ | PRM | 0.79 (0.05) | 0.81 (0.04) | 0.77 (0.05) | 0.82 (0.04) |
|  | SRM |  | 0.99 (0.00) |  | 0.97 (0.01) |
| $CV_{J \rightarrow J}$ | PRM | 0.67 (0.08) | 0.68 (0.07) | 0.62 (0.07) | 0.63 (0.07) |
|  | SRM |  | 0.97 (0.01) |  | 0.96 (0.01) |

Presented is the correlation ( $r$ ) between the predicted mean genetic values and between the predicted specific genetic values. Model 2 comprises a uniform covariance structure for the genetic values while model 4 comprises a compound symmetric covariance structure. IND - *indica*; JAP - *japonica*; PRM - pedigree-based relationship matrix; SRM - site-based relationship matrix; BRM - branch-based relationship matrix. $CV_{IJ \rightarrow IJ}$ trained on and predicted both *indica* and *japonica*, $CV_{I \rightarrow J}$ trained on *indica* and predicted *japonica*, $CV_{J \rightarrow I}$ trained on *japonica* and predicted *indica*, $CV_{I \rightarrow I}$ trained on and predicted *indica*, and $CV_{J \rightarrow J}$ trained on and predicted *japonica*.

Training in both subspecies together gave similar predictive abilities to training within a single subspecies only (Table 4). Models with the BRM had the highest predictive ability for *CV*_*IJ* →*I J*_ and for the within subspecies scenarios *CV*_*I*→ *I*_ and *CV*_*J*→ *J*_. Overall, the predictive abilities were higher for *indica* than *japonica*. The highest predictive abilities (≥0.4) were achieved in *CV*_*IJ* →*I J*_, *CV*_*I* →*I*_, and *CV*_*J* →*J*_ for all models and RMs. Conversely, the lowest predictive abilities were observed for the challenging scenario *CV*_*I* →*J*_, but not for *CV*_*J* →*I*_ where high predictive abilities were achieved with the SRM and BRM, although negative predictive abilities were observed with the PRM. For training in one subspecies and predicting into the other (*CV*_*I*→ *J*_ and *CV*_*J*→ *I*_), there was no consistently best RM. For training and predicting within the same subspecies (*CV*_*I*→ *I*_ and *CV*_*J*→ *J*_), the BRM gave the highest predictive abilities. Overall, the PRM performed unexpectedly well except for *CV*_*J*→ *I*_. The BRM built with *Ne* of 23, 1500, and 150,000 were compared for a favorable scenario (*CV*_*IJ* →*I J*_) and an unfavorable scenario (*CV*_*I*→ *J*_), and predictive abilities with *Ne* = 1500 and 150,000 were the highest and very similar (Supplementary Table 7).

**Table 4.**
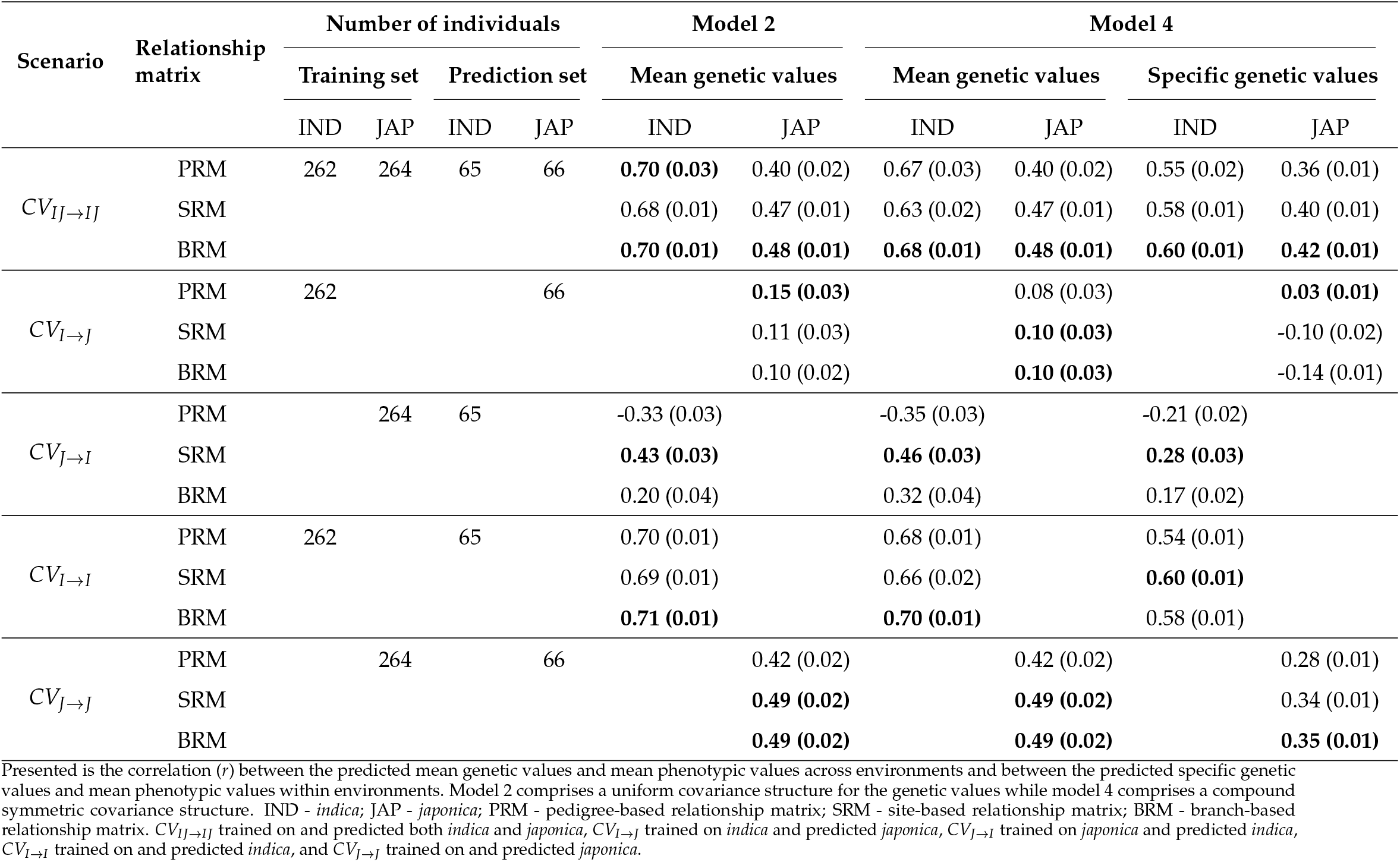
Predictive ability of models 2 and 4 for different relationship matrices and cross-validation scenarios. Presented is the mean predictive ability across 1000 replicates of each scenario, with standard deviations in parentheses. The best-performing combinations are presented in boldface.

The predicted SNP site effects from models 2 and 4 were highly correlated between the SRM and BRM (Table 5). The number of significant SNP site effects for each model and RM was 13 for model 2 and 14 for model 4 with the SRM and 10 for model 2 and 5 for model 4 with the BRM (Figure 6). The standardised SNP site effects for these models are presented in Supplementary Figure 6. A total of 27 SNP site effects were significant across models and RMs. No SNP site effect was significant across all model and RM combinations. Specifically, one SNP site effect was significant across 3 combinations, 13 SNP site effects were significant across two combinations, and the remaining 13 were unique to a single combination.

**Table 5.** Correlation between predicted SNP site effects from models 2 and 4 with the site-based (SRM) and branch-based (BRM) relationship matrices.

| Model | Relationship Matrix | Model 2 |  | Model 4 |  |
| --- | --- | --- | --- | --- | --- |
|  |  | SRM | BRM | SRM | BRM |
| 2 | SRM |  | 0.94 | 0.96 | 0.90 |
|  | BRM | 0.94 |  | 0.91 | 0.97 |
| 4 | SRM | 0.96 | 0.91 |  | 0.94 |
|  | BRM | 0.90 | 0.97 | 0.94 |  |

**Figure 6.**
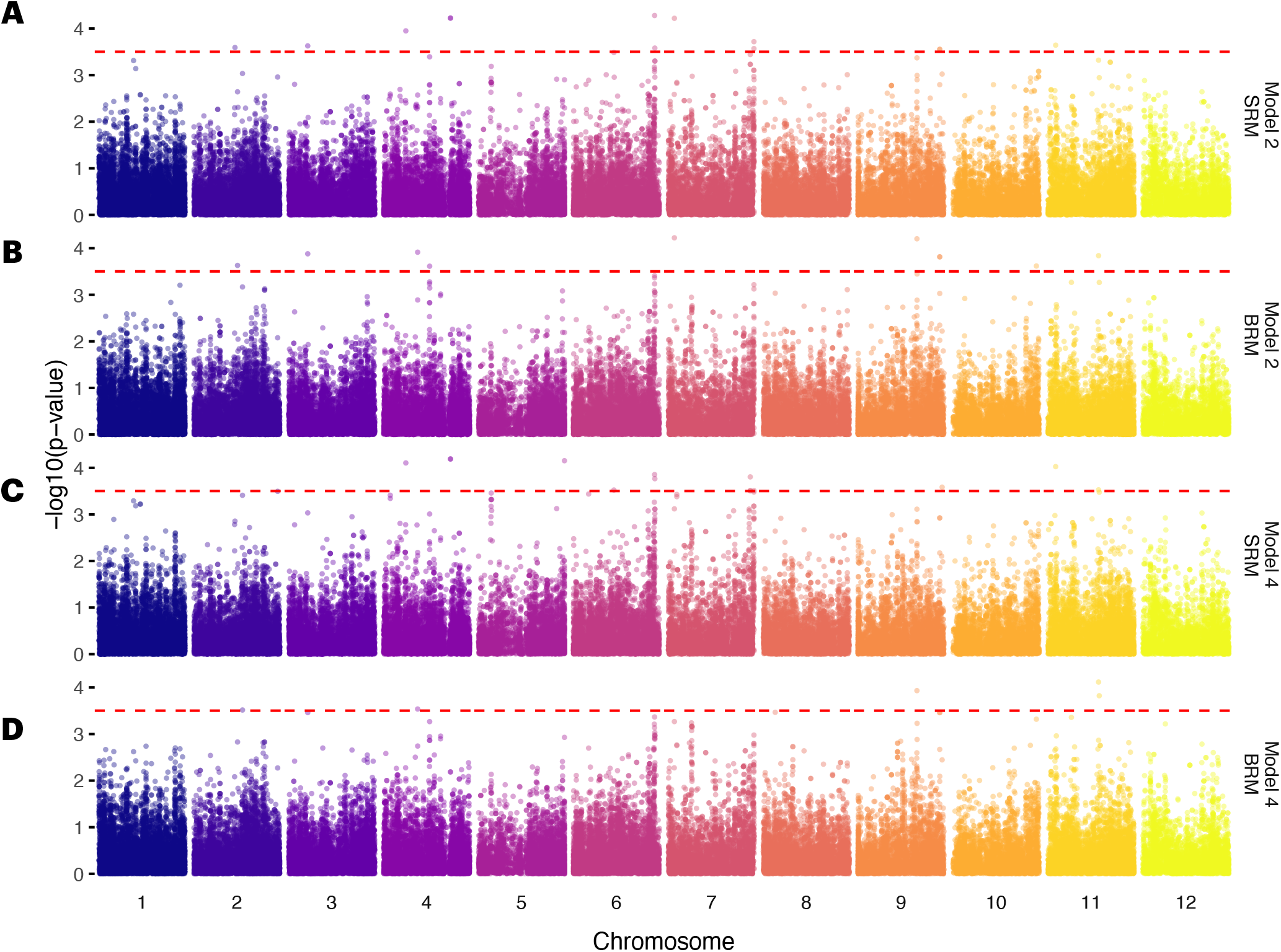
Log-transformed *p*-values for the predicted mean SNP site effects: (A) model 2 with the site-based relationship matrix (SRM); (B) model 2 with the branch-based relationship matrix (BRM); (C) model 4 with the SRM; and (D) Model 4 with the BRM. Model 2 comprises a uniform covariance structure for the genetic values while model 4 comprises a compound symmetric covariance structure. The number of significant SNP site effects was 13 for model 2 and 14 for model 4 with the SRM and 10 for model 2 and 5 for model 4 with the BRM. No SNP site effect was significant across all models and RMs.

We used the zoom-in region to study how the estimated SNP site effects map onto ARG branches and generate haplotype effects. We studied the region between GWAS hits at positions 29,476,724 and 29,561,694 of chromosome 6, which spans 84,970 bp, corresponding to ∼0.3cM. We observed 15 SNP sites in 8 local trees within this region (Supplementary Table 4). Of the 8 trees, 2 contained 4 SNP sites each, 1 tree contained 2 SNP sites, and the remaining 5 contained 1 SNP each. The frequency of SNP mutations was generally higher in *indica* than in *japonica*. The standardised SNP site effect was between -3.55 and 3.52 (Supplementary Table 4). There were a total of 61 haplotypes in the region. The haplotype effect of each local tree ranged from -3.55 to 5.77 for the trees with 4 SNP sites, from 0.00 to 2.58 for the tree with 2 SNP sites, and from -2.78 to 3.28 for the trees with 1 SNP (Supplementary Table 5). Finally, the absolute value of the haplotype effect of the 10 most frequent haplotypes ranged from 0.00 to 3.93 (Supplementary Table 6).

## Discussion

This study adds to the growing body of literature on quantitative genetic analysis across populations (Martin *et al*. 2017; Ding *et al*. 2023; Mester *et al*. 2023; Warburton *et al*. 2023; Zhang *et al*. 2023b) and the applications and benefits of ARGs (Kelleher *et al*. 2019; Wohns *et al*. 2022; Fan *et al*. 2022; Brandt *et al*. 2022; Link *et al*. 2023; Zhang *et al*. 2023a; Brandt *et al*. 2024; Lewanski *et al*. 2024; Sun *et al*. 2024; Gunnarsson *et al*. 2024; Zhu *et al*. 2024; Nielsen *et al*. 2024). We evaluated the use of an ARG to capture the genetic history and improve quantitative genetic modelling in a rice breeding dataset with *indica* and *japonica* subspecies. In the following, we discuss the usefulness of an ARG encoded as a tree sequence for capturing biological features of the analyzed population and for efficiently storing genomic data. We also discuss how tree sequences can capture relationships between the genomes of individuals, making them highly relevant to genomic prediction and GWAS. Finally, we discuss the limitations of the study, particularly the challenges with inferring ARGs and the impact of estimating *Ne* on dating. Overall, our results highlight the potential of ARGs as an emerging tool for quantitative genetic analyses and their application in selective breeding.

### Tree sequences for capturing biological signal and data compression

We corroborated that tree sequences are useful in portraying population structure and are efficient in compressing genomic data. Our GNN analysis successfully captured the underlying population structure and corresponding biological signals; clearly separating the *indica* and *japonica* rice subspecies. The clear within-subspecies structure can be explained by the rice breeding program, which uses a few key individuals as parents for many crosses. The GNN analysis also proved useful for identifying mislabeled individuals and determining the sub-species of unlabelled individuals, increasing their usefulness in future genetic analyses. We found differential local structure by inspecting individual trees in two genome regions of interest concordant with previous studies (Bai *et al*. 2016). Although there were only two SNP sites in each region, we observed a deep split at the *DST* locus between *indica* and *japonica* with a low frequency of both mutations in *indica*. Conversely, we observed that the locus associated with NSB and NSSB segregated in both subspecies. Although there is insufficient data to conclude whether the genomic differentiation observed in the trees is associated with any phenotypic differentiation, the analyses performed here provide greater understanding of the origin and evolution of known QTLs. Such genealogical insight can also aid in differentiating selection and causality from demography responsible for spurious GWAS hits.

The tree sequence format has been shown to be extremely beneficial for dense and whole-genome sequence data (Kelleher *et al*. 2019), and here it compressed the data almost 4 times more than a standard VCF file. Despite this, further work is needed to improve ARG inference in the rice dataset. For example, we inferred 1,031,672 mutations across 61,260 SNP sites, producing an average of ∼17 mutations per site, which is an unexpected result. We followed the approach in Wohns *et al*. (2022), who observed 5,773,816 mutations across 2,090,401 variable SNP sites, producing an average of ∼ 3 mutations per site. The key difference is that they used whole-genome sequencing data, whereas we used reduced-representation GBS data. The high number of mutations per site here may be attributed to the chosen ARG inference parameters; sequencing, imputation and phasing errors; insufficient number of SNP sites; errors in the inferred ancestral allele; or the ARG inference method. While some imputation and phasing errors are expected, the rice individuals are highly inbred so this an unlikely issue, even if the default parameters in Beagle are not optimal for inbred crops such as rice (Niehoff *et al*. 2022).

An alternative inference method may overcome the issues above, e.g., the infinite-sites model, which allows only one mutation per site and thus produces a higher number of inferred recombination events and local trees (Wohns *et al*. 2022; Kelleher *et al*. 2019). Further research is also warranted to optimise the designation of mutation and recombination events (Wohns *et al*. 2022). Such research could use ARGneedle (Zhang *et al*. 2023a) instead of tsinfer (Kelleher *et al*. 2019, 2021b), to also address SNP site ascertainment bias.

### Tree sequences for quantitative genetic analyses

We showed that tree sequences are useful for capturing relationships between the genomes of individuals and can be utilised for genomic prediction and GWAS. The SRM and BRM revealed very similar underlying population structure and were highly correlated, as expected given the duality between these two statistics (Ralph 2019; Ralph *et al*. 2020). Consequently, we observed a high correlation between the predicted mean genetic values obtained with the SRM and BRM for all CV scenarios, particularly when the subspecies in the training and prediction sets were the same. The equivalent correlations involving the PRM were much lower, and even negative when the subspecies in the training and prediction sets were different. This is because predicting from *indica* to *japonica* and vice-versa is very difficult with the PRM due to the lack of deep pedigree and the large number of meiosis events between the subspecies. It is worth noting, however, that the correlations involving the PRM were higher with the BRM than with the SRM for all CV scenarios. This may be attributed to the fact that the BRM and PRM both capture identity by descent information; albeit the BRM captures realised identity by descent from the roots of an ARG and the PRM captures expected identity by descent from the pedigree founders.

All models achieved a high predictive ability when the sub-species in the training and prediction sets were the same. This is expected since the accuracy of genomic prediction is a function of genetic distance between training and prediction individuals (Hickey *et al*. 2014; Lorenz and Smith 2015; Scutari *et al*. 2016; Ding *et al*. 2023). In general, we observed that the BRM achieved the highest predictive ability for these scenarios, highlighting the benefit of ARGs in jointly modelling identity by descent and identity by state relationships. Modest improvements were observed over the SRM, likely due to the limited dataset size, much larger number of parameters estimated (693,524 ARG edges compared to 61,260 SNP sites), and potential errors arising in the underlying ARG.

Training in both subspecies together gave similar predictive abilities to training within a single subspecies only. This is contrary to a previous study involving a subset of our data Berro *et al*. (2019), and could be explained by the larger number of SNP sites used here (61,260 SNP sites compared to 15,545). We observed that *indica* had higher predictive ability than *japonica* for these scenarios, which is expected due to large groups of *indica* relatives within the breeding program (Hickey *et al*. 2014; Scutari *et al*. 2016; Ding *et al*. 2023). Finally, training in one sub-species and predicting into the other subspecies produced the lowest predictive abilities of all scenarios. This is in agreement with previous empirical studies that reported limited predictive ability between populations, let alone between different subspecies like *indica* and *japonica* rice. The results may therefore suggest the presence of a different genetic architecture for complex traits between the subspecies (Zhao *et al*. 2011), different linkage-disequilibrium between typed loci and QTL, or additional (non-additive) genetic effects.

We observed correspondence between the predicted SNP site effects obtained with the SRM and BRM, indicating consistency across models and RMs. Specifically, the significant SNP site effects were spread across different model and RM combinations. Approximately half of the significant SNP site effects were unique to specific model and RM combinations, while the other half were shared across two combinations. Focusing on a specific region of chromosome 6 between two GWAS hits provided additional insights into haplotype effects, revealing variation in SNP and haplotype effects across trees. Such insights will be enhanced in future studies by the presence of known relationship information between SNP sites, allowing unique predictions of site and haplotype effects to be obtained.

### Considerations

Plant breeding datasets are challenged by intense selection from one cycle to the next. We hypothesize that a forward validation scheme could highlight the benefit of more precisely capturing population and family structure with ARGs compared to the five-fold CV used here. Furthermore, the size and population structure of our dataset may be the limiting factor as to why the genomic RMs did not produce much higher predictive abilities than the PRM for the more favourable scenarios *CV*_*IJ*→ *I J*_, *CV*_*I*→ *I*_, and *CV*_*J* → *J*_ (Daetwyler *et al*. 2010; Lorenz and Smith 2015; Papin *et al*. 2024). However, a large portion of the individuals in the dataset did not have genomic data, limiting the study to only those with genomic data available. For these cases, the integration of pedigree and genomic data has been proven successful (Aguilar *et al*. 2010), but future work is needed to integrate them in a single-step version using the BRM.

Depending on the method and software implementation, ARG inference and dating require a high density of SNP sites, knowledge about Ne, ancestral alleles, and mutation and recombination rate. Although rice’s mutation and recombination rate estimates are available (Yang *et al*. 2015; Si *et al*. 2015), the ancestral alleles and *Ne* for our population were not. For 25% of the SNP sites, we were not able to infer the ancestral allele and assumed the major allele as the ancestral. This likely introduced errors, since the chosen ancestral allele was likely to be driven by the higher number of *japonica* individuals in the combined dataset.

The choice of *Ne* is not trivial in agricultural species, since it changes significantly over time due to domestication, bottlenecks, and selective breeding. For rice, we found very contrasting estimates related to different time periods; current 23, recent 1500, and ancient 150,000. The *Ne* assumption strongly impacted the dating of trees, but not the predictive ability of the different models. Finally, we did not use high-density whole-genome sequence data but instead used GBS data with 61,260 SNP sites which can limit the ARG inference. Note, however, it has been shown that accurate ARG inference can be achieved for subsets of whole-genome sequence data, such as SNP arrays, when Ne is small (Fan *et al*. 2022).

### Future directions

There are many new opportunities to expand this research. Firstly, more research is warranted to improve the inference and dating of ARGs, and to understand the inference behaviour for multiple species. Secondly, large-scale datasets are required to achieve sufficient power to estimate the very large number of edge effects, and efficient algorithms are currently underway to compute the BRM for such datasets (P. Ralph, personal communication, August 30th, 2024). Finally, ARGs have the potential to improve genomic prediction and GWAS across diverse populations as they can capture mutations at untyped SNP sites on the inferred edges between genomes (Selle *et al*. 2021; Harris 2023; Link *et al*. 2023; Zhang *et al*. 2023a; Brandt *et al*. 2024; Lewanski *et al*. 2024; Gunnarsson *et al*. 2024; Zhu *et al*. 2024), and may also capture novel epistatic variations responsible for changing mutation effects over time (Mathieson 2021; Park *et al*. 2022).

## Conclusions

We have shown that: (i) an ARG encoded as a tree sequence effectively captured population structure and underlying biological signal in *indica* and *japonica* subspecies in the Uruguayan National Rice Breeding Program. (ii) Tree sequences can be used for genomic prediction and GWAS, and have great potential in larger datasets by estimating mutation effects in different genetic backgrounds. (iii) In terms of GWAS, inspecting the ancestry of haplotypes and their values provides insight into the history of mutations and resulting haplotype differences. (iv) There are still challenges with accurately inferring ARGs. In conclusion, our results highlight the potential of ARGs as an emerging tool for the quantitative genetic analysis of diverse populations.

## Data availability

The dataset, including phenotypic and genomic data, is publicly available at the Dryad repository (Rebollo *et al*. 2023b) with anonymized line identification. Analysis scripts and data are available at https://github.com/HighlanderLab/irebollo_rice_tree.

## Contributions

IR curated and analyzed the data, prepared the results, drafted, and edited the manuscript. DT guided the statistical analysis, derived and wrote the appendix for backsolving SNP site effects from genetic values, contributed to the interpretation of the results, and edited the manuscript. JO contributed to tree sequences inference, interpretation of results, and edited the manuscript. JR led the engagement with the rice breeding program, contributed to data generation and curation, interpretation of results, and edited the manuscript. GG led the study, contributed to the interpretation of results, and edited the manuscript.

## Acknowledgments

The authors acknowledge INIA’s rice breeding team: Pedro Blanco, Fernando Pérez de Vida, Federico Molina, and former and current field and laboratory staff. We also acknowledge Brieuc Lehmann, Jerome Kelleher, and Peter Ralph for tskit implementation of the branch-based relationship matrix (BRM), and Yan Wong for discussions about tree sequences.

## Appendix

This appendix derives an approach to backsolve for SNP site effects from genetic values parameterised by a site-based (SRM) or branch-based (BRM) relationship matrix. Assume the vector of random genetic values in Eq. 1 are modelled as **u** = **M*α***, where **M** is a centered SNP genotype matrix and ***α*** is a vector of SNP site effects, often referred to as allele substitution effects when parameterised by the SRM. It is assumed that:

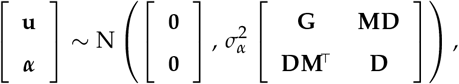

where 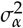 is the variance parameter of the SNP site effects, **G** = **MDM**^⊤^ is the relationship matrix between individuals and **D** is the relationship matrix between SNP site effects. Typically, the matrix **D** is assumed to be diagonal (VanRaden 2008), like for the SRM where **D** = **I**/*c* and *c* is the sum of the expected heterozygosities across SNP sites. In terms of the BRM, however, **D** was unknown and assumed to be non-diagonal, with relationships between sites induced through the tree sequence structure. In this case, we approximated **D**; as:

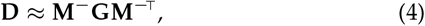

where **M**^−^ is the Moore-Penrose generalised inverse of **M**, obtained such that **D** satisfies **G** ≡ **MDM**^⊤^. We have found that **D** obtained in this manner represents a near-unique solution, and have made brief interpretation in text to reflect this limitation.

Let 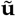 denote the vector of BLUPs of the genetic values parameterised by the site- or branch-based RM. Following Tolhurst *et al*. (2019), the BLUPs of the SNP site effects can be obtained from 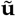 as:

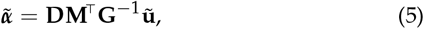

with corresponding variance matrix obtained as:

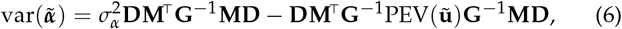

where PEV 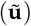 is the prediction error variance matrix of the predicted genetic values, routinely supplied by mixed model software. The test statistics in Eq. 2 are then constructed using the components in Eqs. 5 and 6.

---

**Supplementary Figure 1.**
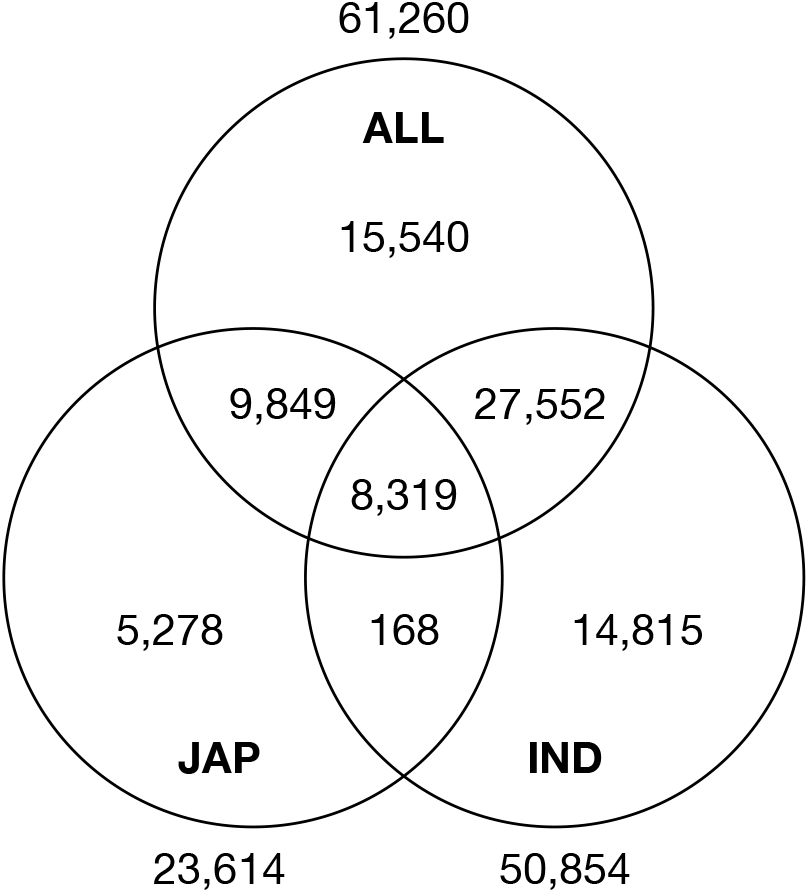
Number of SNP sites unique and shared across the genomic datasets (ALL, both subspecies together; IND, *indica* only; JAP *japonica* only). The number of SNP sites unique to each dataset is given outside the circles, while the number of shared SNP sites is given inside the circles.

**Supplementary Table 1.**
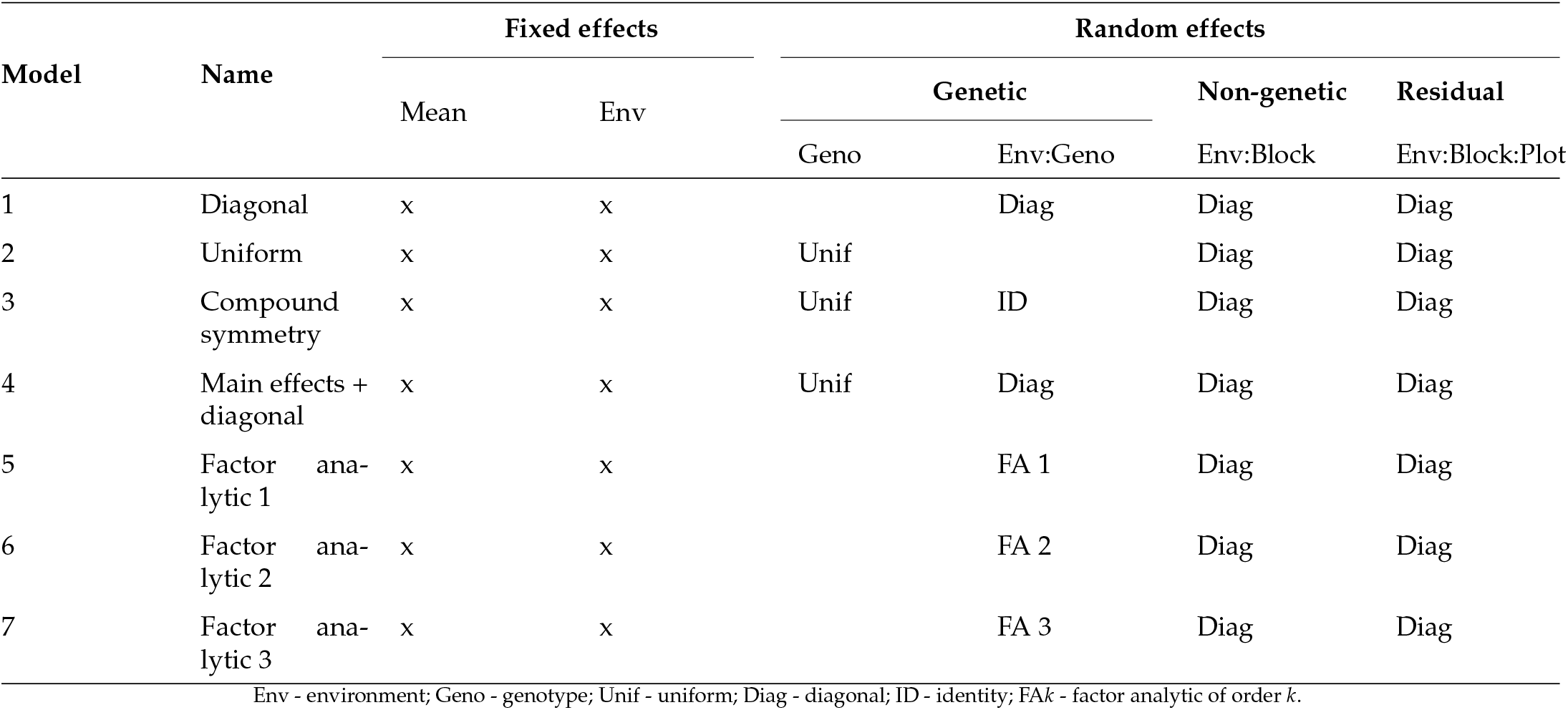
Summary of the fixed effects, random effects, and residual for models 1-7. All models included a fixed overall mean and environment main effects as well as heterogeneous block and residual variances across environments. The models differed in their random genetic terms, as described below.

**Supplementary Table 2.** Residual log-likelihood (LogLik), Akaike Information Criterion (AIC), and percentage of variance explained (*v*_*e*_) for models 1-7 with different relationship matrices.

| Model | LogLik | | | AIC | | | $v_e$ | | |
| --- | --- | --- | --- | --- | --- | --- | --- | --- | --- |
|  | PRM | SRM | BRM | PRM | SRM | BRM | PRM | SRM | BRM |
| 1 | -171,666.2 | -171,527.4 | -171,487.0 | 343,572.5 | 343,294.8 | 343,214.0 |  |  |  |
| 2 | -172,530.8 | -172,408.4 | -172,399.4 | 345,223.6 | 344,978.7 | 344,960.8 |  |  |  |
| 3 | -171,477.2 | -171,324.4 | -171,288.9 | 343,118.4 | 342,812.7 | 342,741.9 | 43.7% | 54.8% | 47.6% |
| 4 | -171,380.8 | -171,240.6 | -171,206.3 | 343,003.6 | 342,723.2 | 342,654.6 | 35.5% | 50.7% | 42.7% |
| 5 | -171,334.7 | -171,202.6 | -171,169.5 | 342,989.4 | 342,725.3 | 342,659.0 | 49.6% | 55.2% | 46.3% |
| 6 | -171,245.4 | -171,145.3 | -171,113.8 | 342,888.8 | 342,688.6 | 342,625.5 | 67.5% | 68.9% | 64.5% |
| 7 | -171,204.9 | -171,145.3 | -171,069.6 | 342,883.8 | 342,707.9 | 342,613.2 | 81.8% | 78.3% | 80.2% |

PRM - pedigree-based relationship matrix; SRM - site-based relationship matrix; BRM - branch-based relationship matrix.

**Supplementary Table 3.** Number of haplotypes in the drought and salt tolerance (*DST*) gene and in the locus associated with the number of panicle secondary branches (NSB) and the number of secondary spikelets per secondary branch (NSSB) for *indica* (IND) and *japonica* (JAP). For both loci, the haplotype was defined as the combination of the alleles at the two SNP sites in the region. The allelic states of each SNP, i.e., ancestral (ANC) or alternative (ALT), are ordered by genomic position and separated by a hyphen. Percentages for each subspecies are presented within parentheses.

|  | Locus |  |  |  |
| --- | --- | --- | --- | --- |
|  | DST |  | NSB & NSSB |  |
|  | IND | JAP | IND | JAP |
| ANC-ANC | 30 (4%) | 1084 (98%) | 208 (27%) | 889 (80%) |
| ANC-ALT | 145 (19%) | 3 (0%) | 542 (71%) | 8 (1%) |
| ALT-ANC | 585 (77%) | 23 (2%) | 8 (1%) | 213 (19%) |
| ALT-ALT | 2 (0%) | 0 (0%) | 4 (1%) | 0 (0%) |

**Supplementary Table 4.** Local trees and their SNP covering the zoom-in region, bordered by two SNP sites found significant in the genome-wide association study (chromosome 6, positions 29476724 to 29561694), and the number of genotypes with alternative (i.e., non-ancestral) state in *indica* (IND) and *japonica* (JAP) and corresponding percentage for each subspecies within parentheses, along with the standardised SNP site effect (*z*_*i*_).

| Local tree | SNP | IND | JAP | $z_i$ |
| --- | --- | --- | --- | --- |
| Tree 1 | S6_29476724 | 8 (1%) | 366 (33%) | +3.44 |
|  | S6_29476748 | 175 (23%) | 66 (6%) | +0.35 |
|  | S6_29476763 | 530 (70%) | 399 (36%) | +3.52 |
|  | S6_29476787 | 531 (70%) | 396 (36%) | -3.55 |
| Tree 2 | S6_29480408 | 535 (70%) | 400 (36%) | -2.78 |
| tree 3 | S6_29480471 | 530 (70%) | 397 (36%) | +2.58 |
|  | S6_29480530 | 529 (69%) | 397 (36%) | +2.53 |
| Tree 4 | S6_29516992 | 520 (68%) | 27 (2%) | -0.70 |
| Tree 5 | S6_29517004 | 5 (1%) | 357 (32%) | +2.17 |
| Tree 6 | S6_29531546 | 753 (99%) | 457 (41%) | +3.03 |
| Tree 7 | S6_29557666 | 751 (99%) | 453 (41%) | +3.11 |
|  | S6_29557756 | 738 (97%) | 99 (9%) | +0.55 |
|  | S6_29557803 | 13 (2%) | 352 (32%) | +2.28 |
|  | S6_29561680 | 751 (99%) | 456 (41%) | +3.20 |
| Tree 8 | S6_29561694 | 751 (99%) | 409 (37%) | +3.28 |

**Supplementary Figure 2.**
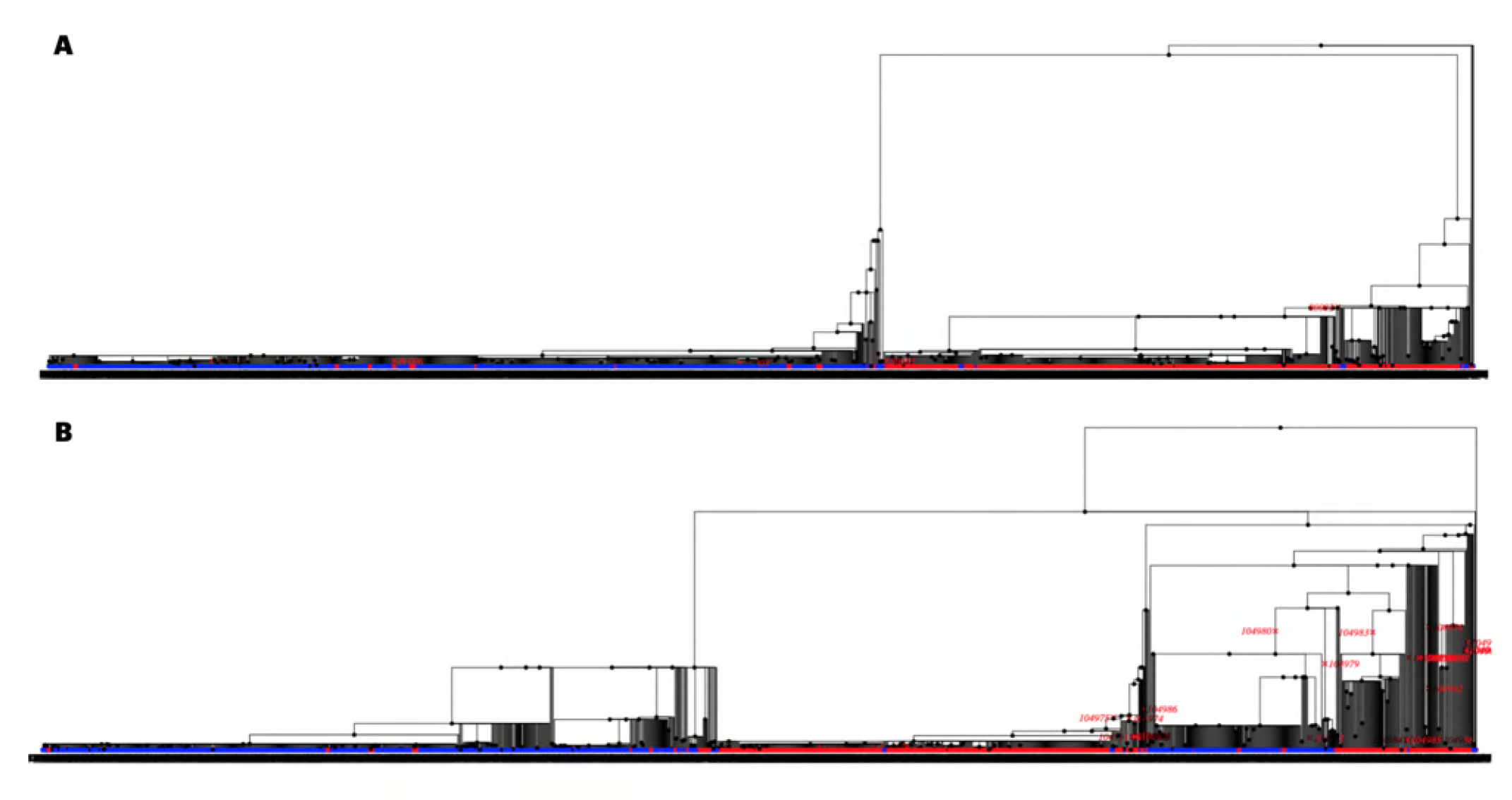
Differential local tree structure at the ending SNP covering the genome positions in (A) the Drought and Salt Tolerance (*DST*) gene, associated with panicle length only in *japonica* and (B) a locus associated with number of panicle secondary branches (NSB) and number of secondary spikelets per secondary branch (NSSB) in both *indica* and *japonica*. The tree in (A) shows a very clear and deep separation between *indica* and *japonica* while the tree in (B) shows segregation in both *indica* and *japonica*.

**Supplementary Table 5.** Haplotypes and their number of genotypes in *indica* (IND) and *japonica* (JAP) and corresponding percentage for each subspecies within parentheses, along with the standardised haplotype effect (*h*_*j*_) for each local tree covering the zoom-in region, bordered by two SNP sites found significant in the genome-wide association study (chromosome 6, positions 29,476,724 to 29,561,694). Haplotypes were defined as the combination of the allelic state (0 for ancestral and 1 for alternative) of all SNP sites found in each tree, ordered by genomic position.

| Local tree | Haplotype | IND | JAP | $h_j$ |
| --- | --- | --- | --- | --- |
| Tree 1 Chr6:<br>29,476,724 -<br>29,476,787 | 0000 | 55 (7%) | 645 (58%) | 0.00 |
|  | 0001 | 2 (0%) | 0 (0%) | -3.55 |
|  | 0010 | 3 (1%) | 1 (0%) | +3.52 |
|  | 0011 | 519 (68%) | 32 (3%) | -0.03 |
|  | 0100 | 173 (23%) | 66 (6%) | +0.35 |
|  | 0101 | 2 (0%) | 0 (0%) | -3.04 |
|  | 1010 | 0 (0%) | 2 (0%) | +3.83 |
|  | 1011 | 8 (1%) | 364 (33%) | +1.21 |
| Tree 2 Chr6:<br>29,480,408 | 0 | 227 (30%) | 710 (64%) | 0.00 |
|  | 1 | 535 (70%) | 400 (36%) | -2.78 |
| Tree 3 Chr6:<br>29,480,471 -<br>29,480,530 | 00 | 232 (30%) | 712 (64%) | 0.00 |
|  | 01 | 0 (0%) | 1 (0%) | +2.53 |
|  | 10 | 1 (0%) | 1 (0%) | +2.58 |
|  | 11 | 529 (70%) | 396 (36%) | +2.56 |
| Tree 4 Chr6:<br>29,516,992 | 0 | 242 (32%) | 1083 (98%) | 0.00 |
|  | 1 | 520 (68%) | 27 (2%) | -0.7 |
| Tree 5 Chr6:<br>29,517,004 | 0 | 757 (99%) | 753 (68%) | 0.00 |
|  | 1 | 5 (1%) | 357 (32%) | +2.17 |
| Tree 6 Chr6:<br>29,531,546 | 0 | 9 (1%) | 653 (59%) | 0.00 |
|  | 1 | 753 (99%) | 457 (41%) | +3.03 |
| Tree 7 Chr6:<br>29,557,666 -<br>29,561,680 | 0000 | 11 (1%) | 652 (59%) | 0.00 |
|  | 0001 | 0 (0%) | 3 (0%) | +3.20 |
|  | 0011 | 0 (0%) | 1 (0%) | +5.77 |
|  | 0100 | 0 (0%) | 1 (0%) | +0.55 |
|  | 1000 | 0 (0%) | 1 (0%) | +3.11 |
|  | 1001 | 0 (0%) | 3 (0%) | +3.27 |
|  | 1011 | 13 (2%) | 351 (32%) | +5.56 |
|  | 1101 | 738 (97%) | 98 (9%) | +3.68 |
| Tree 8 Chr6:<br>29,561,694 | 0 | 11 (1%) | 701 (63%) | 0.00 |
|  | 1 | 751 (99%) | 409 (37%) | +3.28 |

**Supplementary Table 6.**
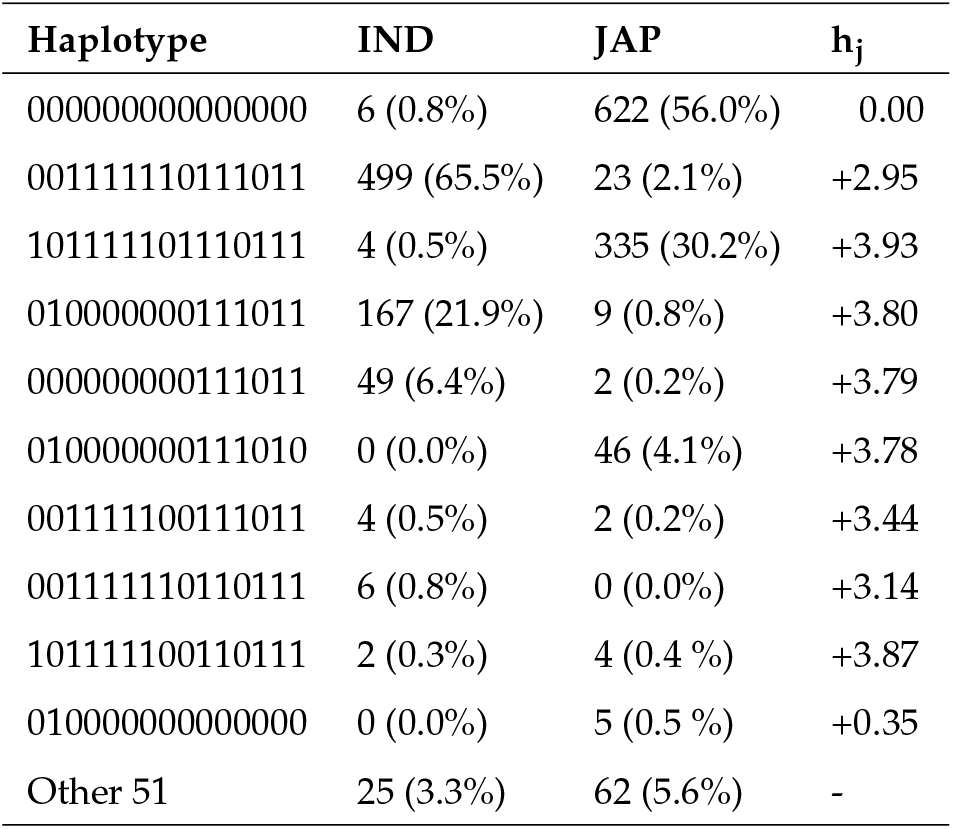
The 10 most frequent haplotypes, their number of genotypes in *indica* (IND) and *japonica* (JAP) and corresponding percentage for each subspecies within parentheses, and their standardised effect (*h*_*j*_) for the zoom-in region, bordered by two SNP sites found significant in the genome-wide association study (chromosome 6, positions 29,476,724 to 29,561,694). Haplotypes were defined as the combination of the allelic state (0 for ancestral and 1 for alternative) of all SNP sites found in the zoom-in region, ordered by genomic position.

**Supplementary Table 7.**
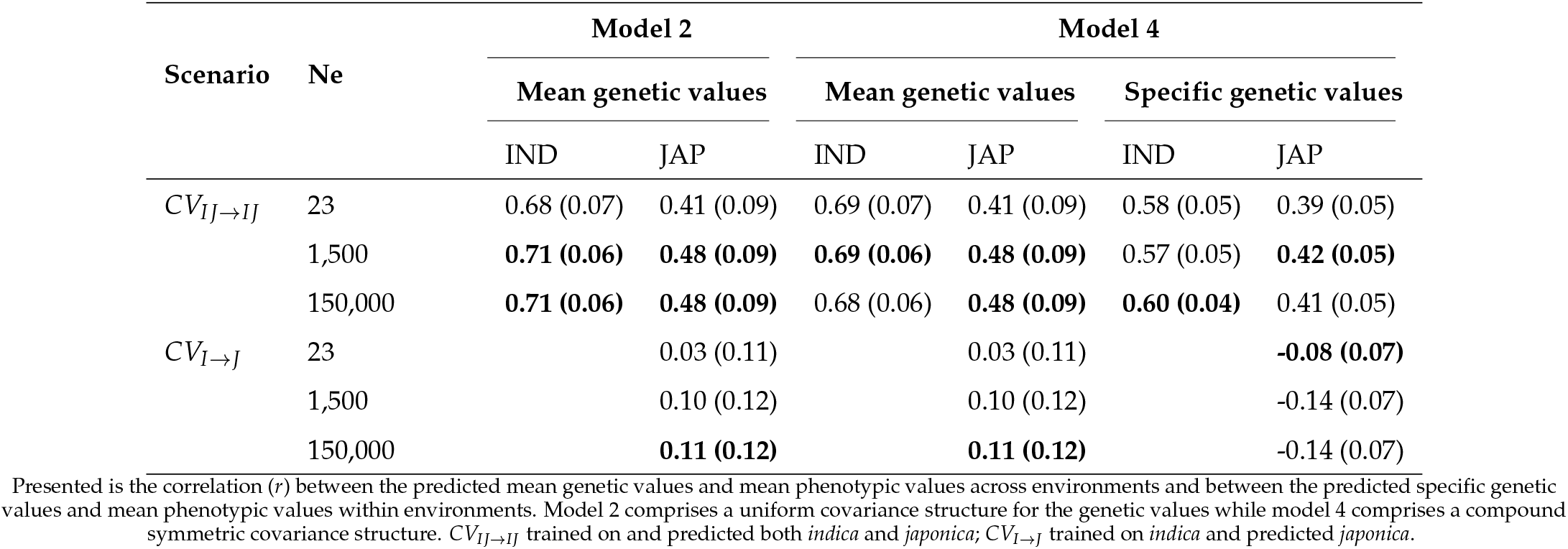
Predictive ability of models 2 and 4 with the branch-based relationship matrix (BRM) built with three effective population sizes (*Ne*) for two cross-validation scenarios in *indica* (IND) and *japonica* (JAP). Values presented are the mean predictive ability across 100 replicates of each scenario, with standard deviations provided in parentheses. Results of the best-performing *Ne* for each scenario and population are presented in boldface.

| Scenario | $N_e$ | Model 2 | | Model 4 | | | |
| --- | --- | --- | --- | --- | --- | --- | --- |
|  |  | Mean genetic values |  | Mean genetic values |  | Specific genetic values |  |
|  |  | IND | JAP | IND | JAP | IND | JAP |
| $CV_{IJ \rightarrow IJ}$ | 23 | 0.68 (0.07) | 0.41 (0.09) | 0.69 (0.07) | 0.41 (0.09) | 0.58 (0.05) | 0.39 (0.05) |
|  | 1,500 | **0.71 (0.06)** | **0.48 (0.09)** | **0.69 (0.06)** | **0.48 (0.09)** | 0.57 (0.05) | **0.42 (0.05)** |
|  | 150,000 | **0.71 (0.06)** | **0.48 (0.09)** | 0.68 (0.06) | **0.48 (0.09)** | **0.60 (0.04)** | 0.41 (0.05) |
| $CV_{I \rightarrow J}$ | 23 | | 0.03 (0.11) | | 0.03 (0.11) | | **-0.08 (0.07)** |
|  | 1,500 |  | 0.10 (0.12) |  | 0.10 (0.12) |  | -0.14 (0.07) |
|  | 150,000 |  | **0.11 (0.12)** |  | **0.11 (0.12)** |  | -0.14 (0.07) |

Presented is the correlation ( $r$ ) between the predicted mean genetic values and mean phenotypic values across environments and between the predicted specific genetic values and mean phenotypic values within environments. Model 2 comprises a uniform covariance structure for the genetic values while model 4 comprises a compound symmetric covariance structure. $CV_{IJ \rightarrow IJ}$ trained on and predicted both *indica* and *japonica*; $CV_{I \rightarrow J}$ trained on *indica* and predicted *japonica*.

**Supplementary Figure 3.**
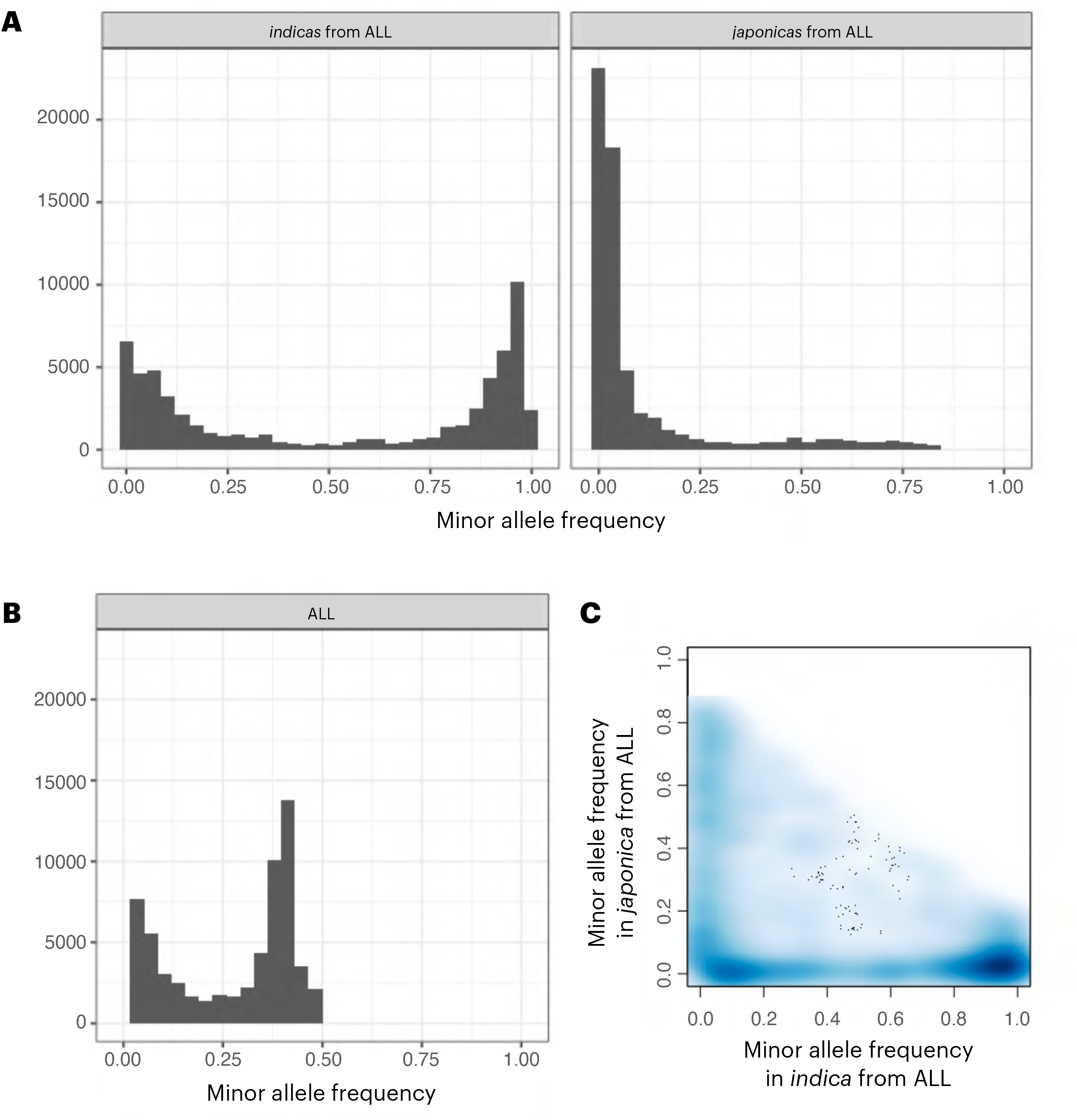
Minor allele frequency spectrum. (A) Minor allele frequency of the subset of *indica* and *japonica* from the ALL genomic dataset that includes all individuals from both subspecies, (B) minor allele frequency of the ALL genomic dataset, and (C) comparison between the minor allele frequency between *indica* and *japonica* from the ALL genomic dataset.

**Supplementary Figure 4.**
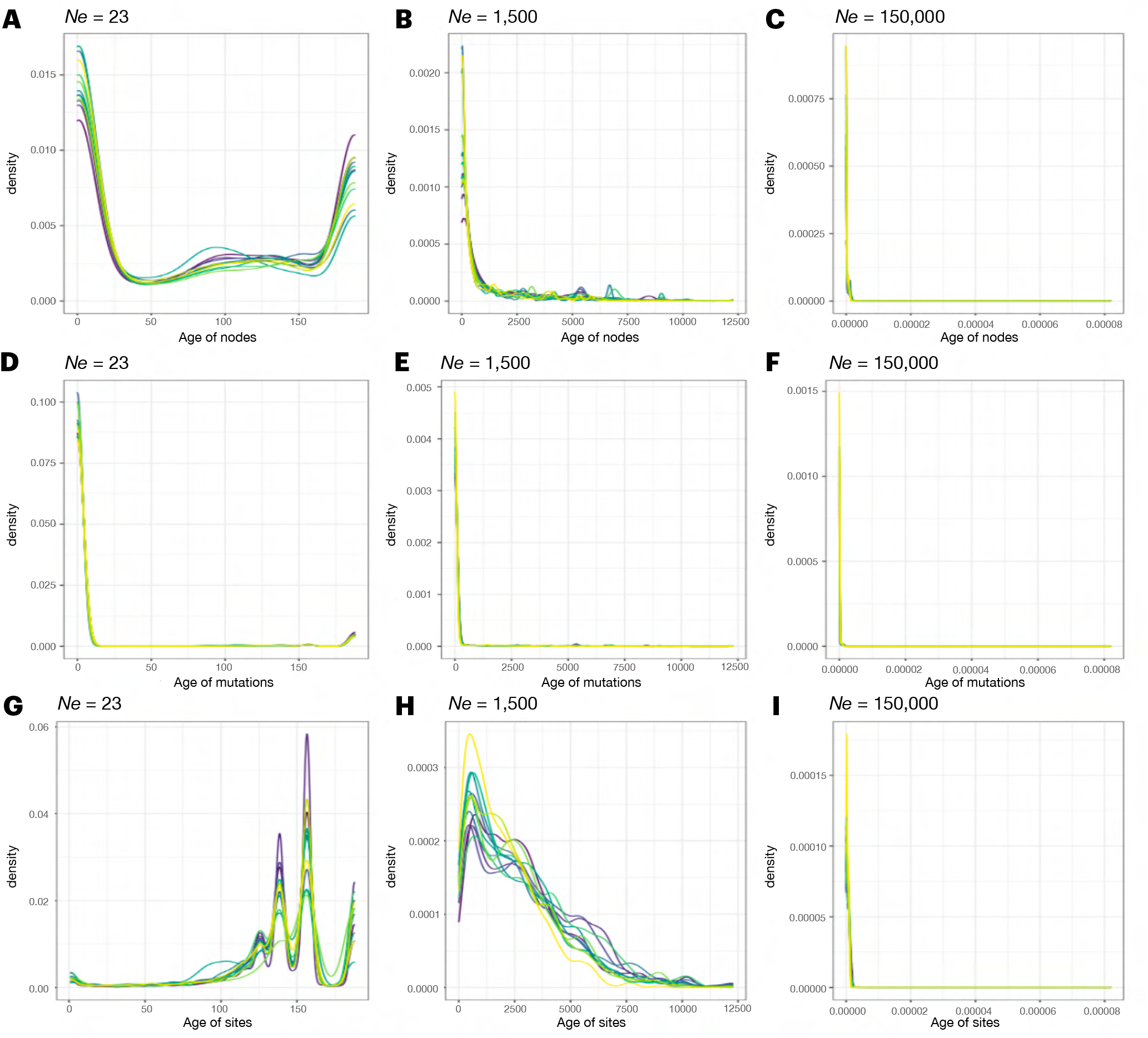
Estimated age of nodes (A, B, and C), mutations (D, E, and F), and sites (G, H, and I) per chromosome with three effective population sizes (*Ne*). The age of nodes represents the age of ancestors in the trees. The age of sites represents the age at which sites became polymorphic, which equals the age of the oldest ancestor above which we inferred a mutation.

**Supplementary Figure 5.**
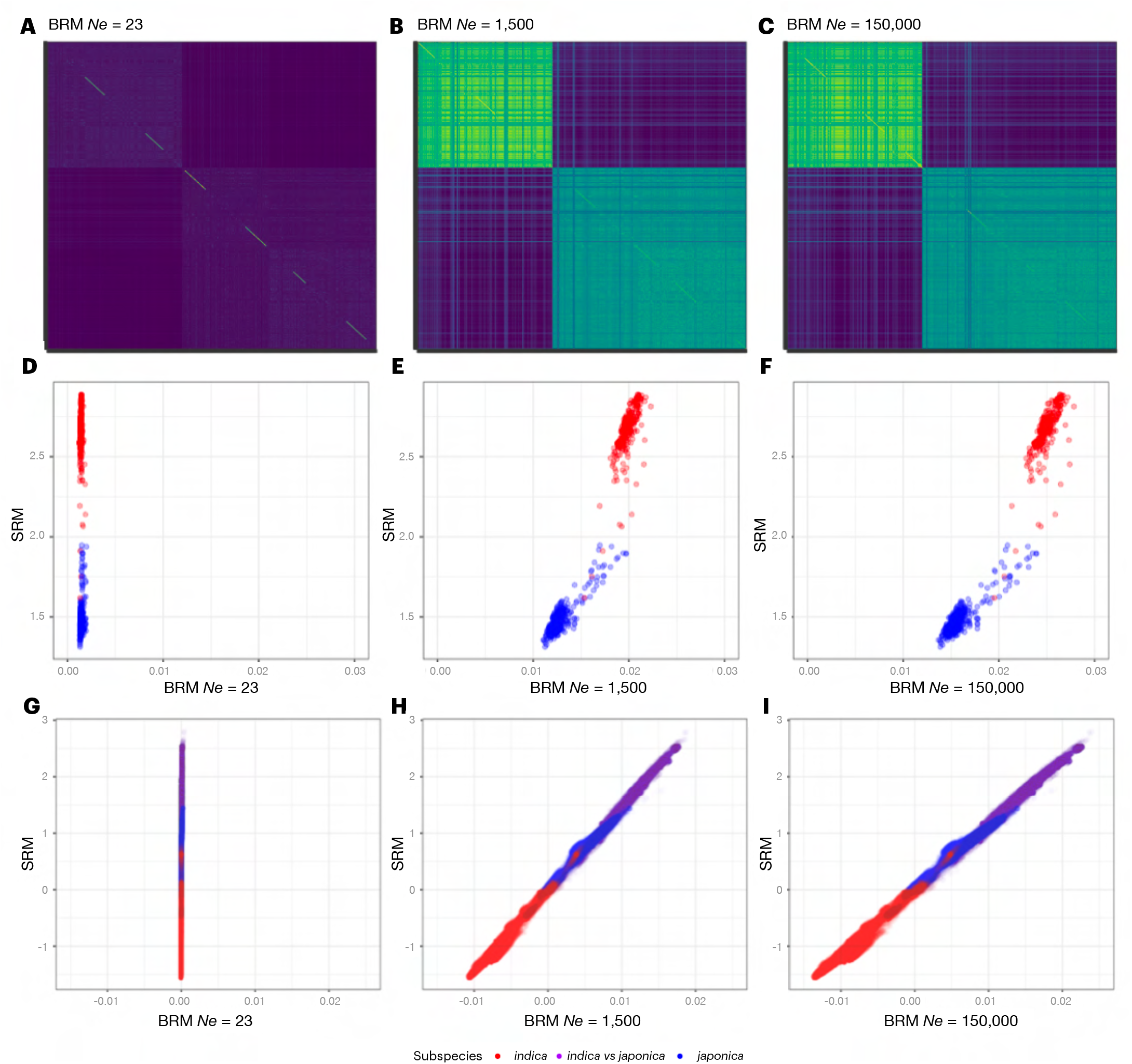
Comparison of the site-based (SRM) and branch-based (BRM) relationship matrices with three effective population sizes (*Ne*). Heatmap of the BRM (A, B, and C), comparison of the diagonal (D, E, and F), and off-diagonal (G, H, and I) elements of both matrices.

**Supplementary Figure 6.**
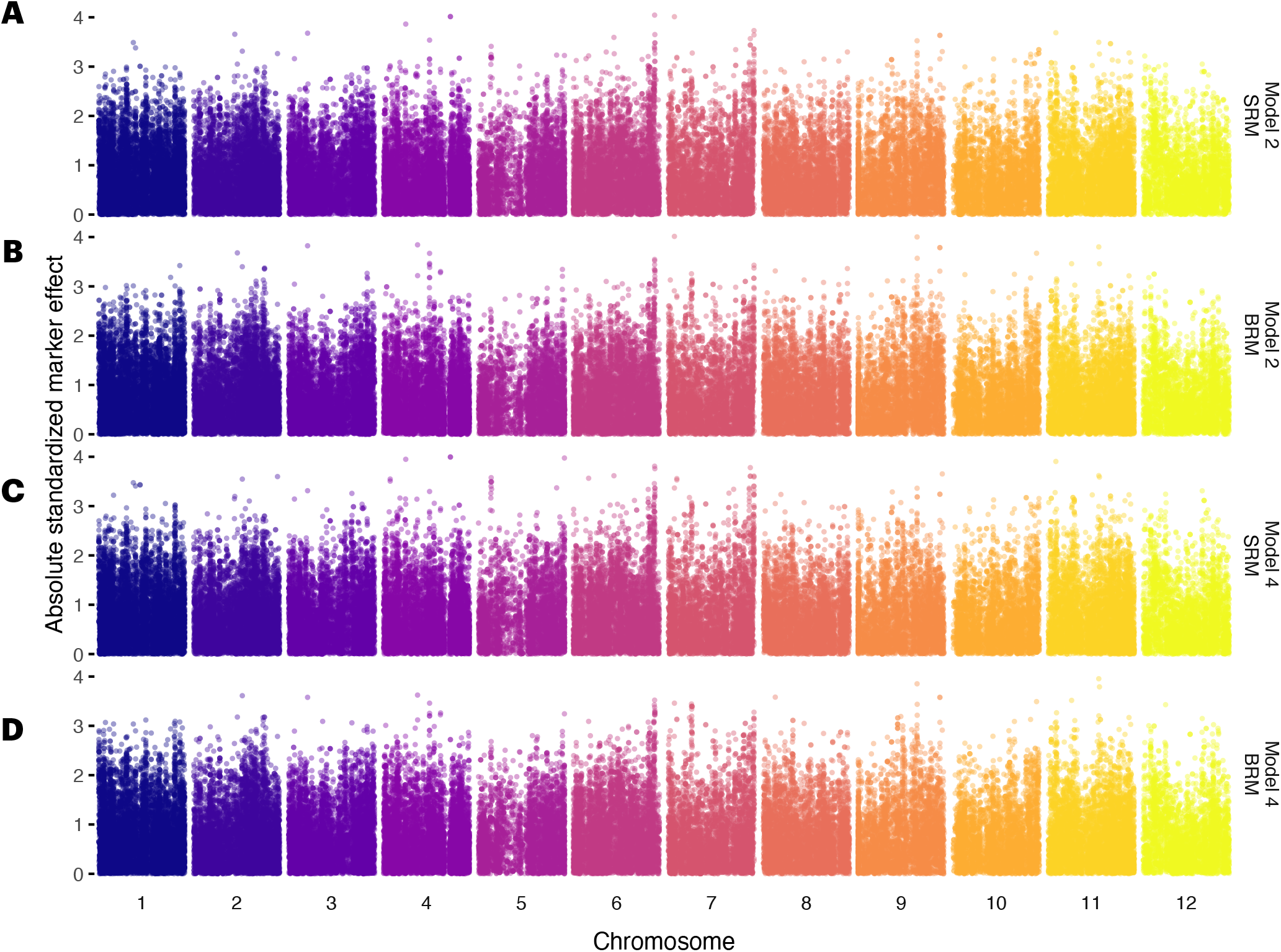
Standardised best linear unbiased predictions of the SNP site effects obtained with the site-based relationship matrix (SRM) from models 2 (A) and 4 (B), and the branch-based relationship matrix (BRM) from models 2 (C) and 4 (D).

